# A non-canonical role and regulations of polo-like kinase-4 in fibroblast cell-type transition

**DOI:** 10.1101/570267

**Authors:** Jing Li, Go Urabe, Mengxue Zhang, Yitao Huang, Bowen Wang, Lynn Marcho, Hongtao Shen, K. Craig Kent, Lian-Wang Guo

**Author notes:** Corresponding author: Lian-Wang Guo, Ph.D., Department of Surgery and Department of Physiology & Cell Biology, The Ohio State University, Columbus, OH 43210, USA.

## Abstract

A divergent member of the polo-like kinase family, PLK4 is known for its canonical role in centriole duplication. Its non-canonical function and regulators are poorly defined. Here we investigated PLK4’s activation and expression and regulations thereof in rat adventitial fibroblast cell-type transition induced by platelet-derived growth factor (PDGF-AA).

Experiments using siRNA and selective inhibitor (centrinone-B) revealed a role for PLK4 not only in AA-induced proliferation/migration, but also in serum response factor (SRF) activation and smooth muscle α-actin expression. PDGFR (receptor) inhibition abrogated AA-stimulated PLK4 activation (phosphorylation) and expression; P38 inhibition (siRNA, inhibitor) downstream of PDGFR also mitigated PLK4 activation. Furthermore, transcription of PLK4 (and PDGFRα) was repressed by pan-inhibition of the bromodomain/extraterminal family of chromatin-bookmark readers (BRD2, BRD3, BRD4), an effect determined herein as mainly mediated by BRD4. In vivo, periadventitial administration of centrinone-B reduced collagen content and thickness of the adventitia in a rat model of carotid artery injury.

In summary, we have identified a non-canonical role for PLK4 in SRF activation and its regulations by BRD4/PDGFRα-dominated pathways. Results in this study suggest PLK4 inhibition as a potential anti-fibrotic intervention.

## Introduction

Polo-like kinases regulate cell cycle entry and exit. Among the five PLK family members, PLK1 has well-documented roles in multiple steps of mitosis. PLK4, the divergent family member with little homology to the other four PLKs^1^, is reportedly a master regulator of centriole duplication^2^, but its non-canonical functions are still obscure^3^. Moreover, while a few substrates of PLK4 kinase activity were recently identified (e.g. STIL in centriole formation), regulators of PLK4 activation and expression are thus far poorly defined^3, 4^. Recent literature implicates PLKs in fibrogenic processes. For example, PLK1 is found to be a target gene of FoxM1, a transcription factor that promotes lung fibrosis^5^’^6^. This raises the question concerning whether fibroblast cell-type transitions involved in fibrosis are regulated by one or more PLKs.

A well-known fibroblast cell-type change is its transition into myofibroblast, a fibrogenic process involved in numerous disease conditions^7^. Although the definition of myofibroblasts is still debated, these cells are generally characterized by smooth muscle-like morphologies and proliferative/ migratory behaviors^8^. Moreover, they often exhibit high levels of smooth muscle α-actin (αSMA), vimentin, platelet derived growth factor receptor α (PDGFRα), and extracellular matrix proteins (e.g. collagen). Whereas a variety of cell types can differentiate into myofibroblasts^7^, resident fibroblasts have been confirmed as the main source in recent in vivo lineage tracing studies, at least in some vital organs such as heart^8, 9^. It is thus important to identify the regulatory mechanisms in fibroblast cell-type transition. This complex process involves extracellular and cell membrane signaling, cytosolic pathways, epigenetic and transcriptional remodeling, and interactions among these networks^8^. The best known fibrogenic signaling pathway is transforming growth factor (TGFβ1). By contrast, the PDGF pathways are less well-understood^8^. In particular, the PDGF-AA homodimer, which selectively activates PDGFRα, is inadequately explored relative to PDGF-BB, which activates both PDGFRα and PDGFRβ^10^.

In this study, we investigated a possible PLK regulation of vascular adventitial fibroblast cell-type transition in the setting of PDGF-AA-stimulated PDGFRα activation. We focused primarily on the divergent PLK member (PLK4)^3^,and also included PLK1, the representative member of the PLK family^11^. We found that PLK4 inhibition constrained the rat aortic fibroblast proliferative/migratory behaviors, and also the nuclear activity of serum response factor (SRF), a master transcription factor^8^. The latter finding is somewhat surprising, given that PLK4 is deemed as centriole-specific and cytosol-localized. We also uncovered that PDGFR and downstream kinase P38 positively regulated PLK4 activation. In pursuit of the transcriptional regulators of PLK4, we identified BRD4 (a bromodomain/extraterminal family member) as an epigenetic determinant of PLK4 expression. Hence, we have identified a non-canonical function of PLK4. Furthermore, we also observed an effect of PLK4 inhibition on attenuating vascular fibrosis in a rat artery injury model.

## Results

### PLK4 inhibition blocks PDGF-AA stimulated cell-type transition of rat aortic adventitial fibroblasts

For the current studies, we needed to reliably induce fibroblast cell-type transition. To do so, we used PDGF-AA (hereafter abbreviated as AA), which preferentially activates PDGFRα *vs* PDGFRβ^10^. We validated AA’s ligand functionality by demonstrating that it mimicked TGFβ1^7^, the best known potent stimulator of αSMA expression in rat aortic adventitial fibroblasts (Figure S1). Treatment of cells with AA induced an elongated morphology and ∼3-fold increases of proliferation and migration, indicative of fibroblast cell-type transition (Figure 1, A-C).

**Figure 1.**
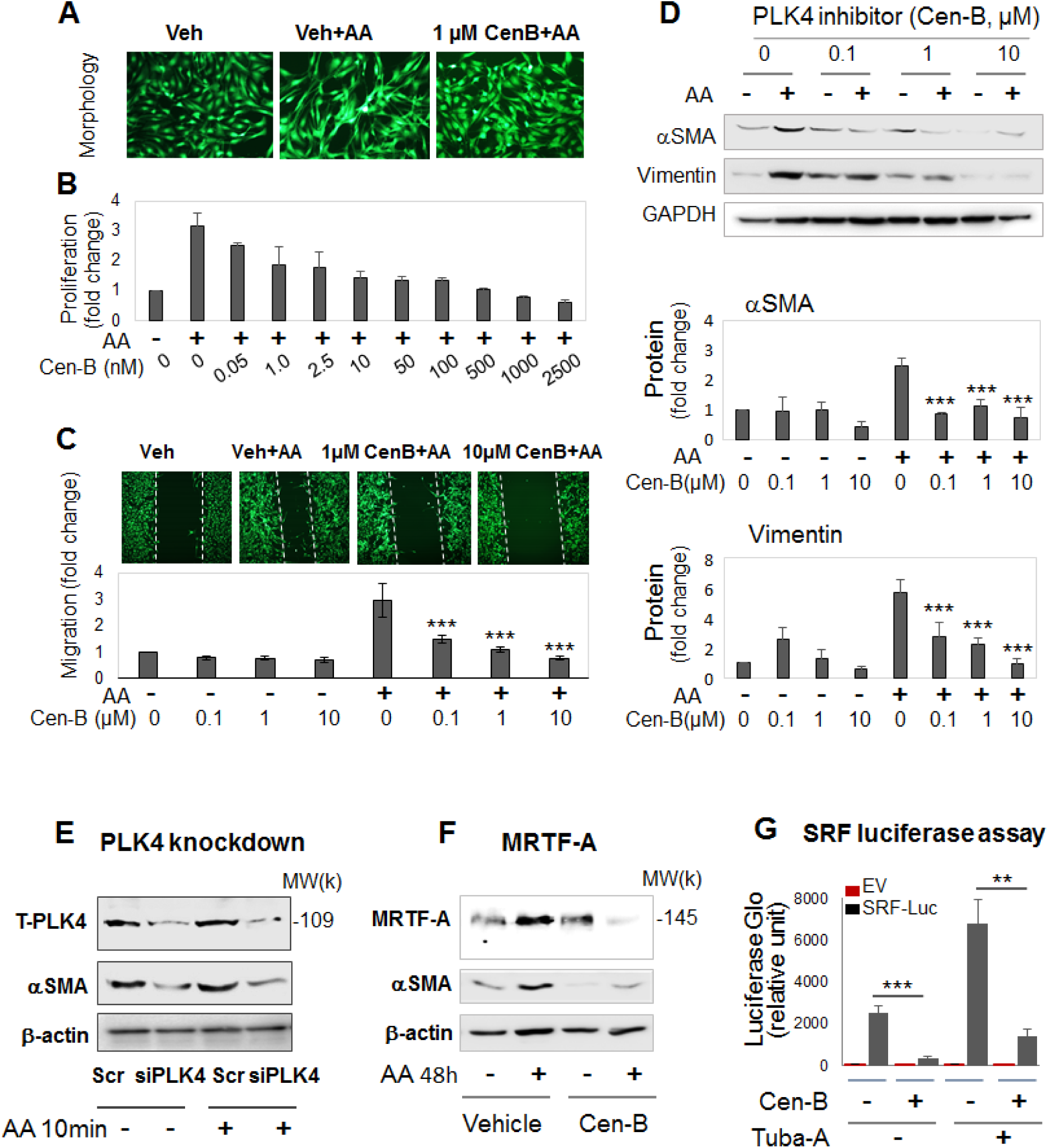
PLK4 inhibition blocks cell-type transition and αSMA expression of rat adventitial fibroblasts. Rat primary adventitial fibroblasts were cultured in the complete medium, starved in the basal medium overnight (see Methods), and pretreated for 2h with vehicle (equal amount of DMSO) or the PLK4-selective inhibitor centrinone-B (Cen-B) at indicated concentrations, followed by stimulation with 60 ng/ml PDGF-AA (abbreviated as AA throughout). Cells were harvested at 24h after stimulation (or otherwise specifically indicated) for various assays. A. Morphology. Cells were (or not) stimulated by AA for 24h without or with pre-treatment (1µM Cen-B). Green fluorescent calcein was used to illuminate cell morphologies. B. Proliferation. CellTiterGlo assay was performed after 72h AA stimulation of cells without or with pretreatment by Cen-B at increasing concentrations. C. Migration (scratch assay). Cells were (or not) stimulated by AA for 24h without or with pre-treatment (1 or 10 µM Cen-B). Calcein was used to illuminate the cells. D. Western blots of αSMA and vimentin. Protein band densitometry was normalized to loading control (β-actin), and then to the basal condition (DMSO, no AA), and finally quantified as fold change. Fold changes from at least 3 independent experiments were averaged, and mean ± SEM was calculated. Quantification for C and D: Mean ± SEM, n ≥3 experiments. One-way ANOVA/Bonferroni test: ***P< 0.001 compared to the condition of AA without Cen-B. E. Effect of PLK4 silencing. Cells were transfected with scrambled or PLK4-specific siRNA for 48 h in the complete medium, starved overnight, and then stimulated with AA for 10 min before harvest for Western blotting. F. Effect of PLK4 inhibition on MRTF-A. Cells were pretreated with 1µM Cen-B (or vehicle) and then stimulated (or not) with AA for 48 h. G. Luciferase reporter assay of SRF transcriptional activity. Cells were transfected with the empty vector control or the SRF-Luciferase vector, followed by luminescence reading. The condition with 5 µM tubastatin-A, an HDAC6 inhibitor and a novel SRF stimulator^12^, served as a positive control. Mean ± SEM, n = 3 experiments. Student’s t-test: **P <0.01, ***P<0.001, between two gray bars.

We then investigated the effect of manipulating PLK4 on this AA-induced process. Here, we took advantage of recent progress in developing highly selective PLK inhibitors that provide powerful tools for deciphering PLK4 functions. Centrinone-B (herein abbreviated as Cen-B) is a novel PLK4-selective inhibitor (Ki = 0.6 nM) with very low affinities for other PLK and non-PLK kinases^1^. Pretreatment of the rat aortic fibroblasts with Cen-B abrogated AA-stimulated proliferation and migration in a concentration-dependent fashion (Figure 1, B and C) (Figure S2). Furthermore, AA-stimulated upregulation of αSMA and vimentin proteins (2.5-5 fold) was also abolished by pretreatment with Cen-B (Figure 1D).

Importantly, we confirmed the PLK4 functional specificity. We did this by silencing PLK4 with siRNA and showing that αSMA levels were reduced (Figure 1E). While it is intuitive that PLK4 as a mitotic factor promoted cell proliferation^1^, a role for PLK4 in elevating αSMA expression was somewhat unexpected given PLK4’s canonical association with centrioles in the cytoplasm.

We then investigated the mechanism underlying PLK4-stimulated αSMA expression. αSMA transcription is known to be driven by SRF, a master transcription factor. SRF is itself activated by MRTF-A, a powerful transcription regulator that shuttles between the cytoplasm and nucleus^12^. We therefore investigated the influence of PLK4 on MRTF-A protein and SRF transcriptional activity. We found that, while treatment with AA elevated MRTF-A protein levels, PLK4 inhibition with Cen-B prevented this elevation (Figure 1F). Furthermore, the SRF transcriptional (luciferase) activity was diminished by Cen-B (Figure 1G). These results revealed that, in rat aortic adventitial fibroblasts, PLK4 promotes SRF activation and αSMA production, at least in part by elevating MRTF-A protein levels.

Taken together, our results indicate that PLK4 regulates fibroblast cell-type transition, and intriguingly, also SRF nuclear activity, a function that apparently departs from that in centriole duplication. To the best of our knowledge, this non-canonical PLK4 function in SRF activation was not previously reported. It is also noteworthy that pretreatment with Cen-B largely preserved normal fibroblastic phenotypes (Figure 1, A-D) and did not cause obvious cell death even at high (e.g.10 μM) concentrations, suggesting a low cytotoxicity of this drug.

We also determined the effect of PLK1 inhibition on fibroblast phenotypes using the PLK1-selective inhibitor GSK461364 (Ki = 2.2 nM, hereafter abbreviated as G-4)^13^. The result (Figure 2) (Figure S3) was similar to that of PLK4 inhibition. However, the PLK1 inhibitor at high concentrations (e.g. 0.5 μM) reduced cell viability to much below the non-stimulated basal level (Figure 2C), consistent with its cytotoxicity^13^.

**Figure 2.**
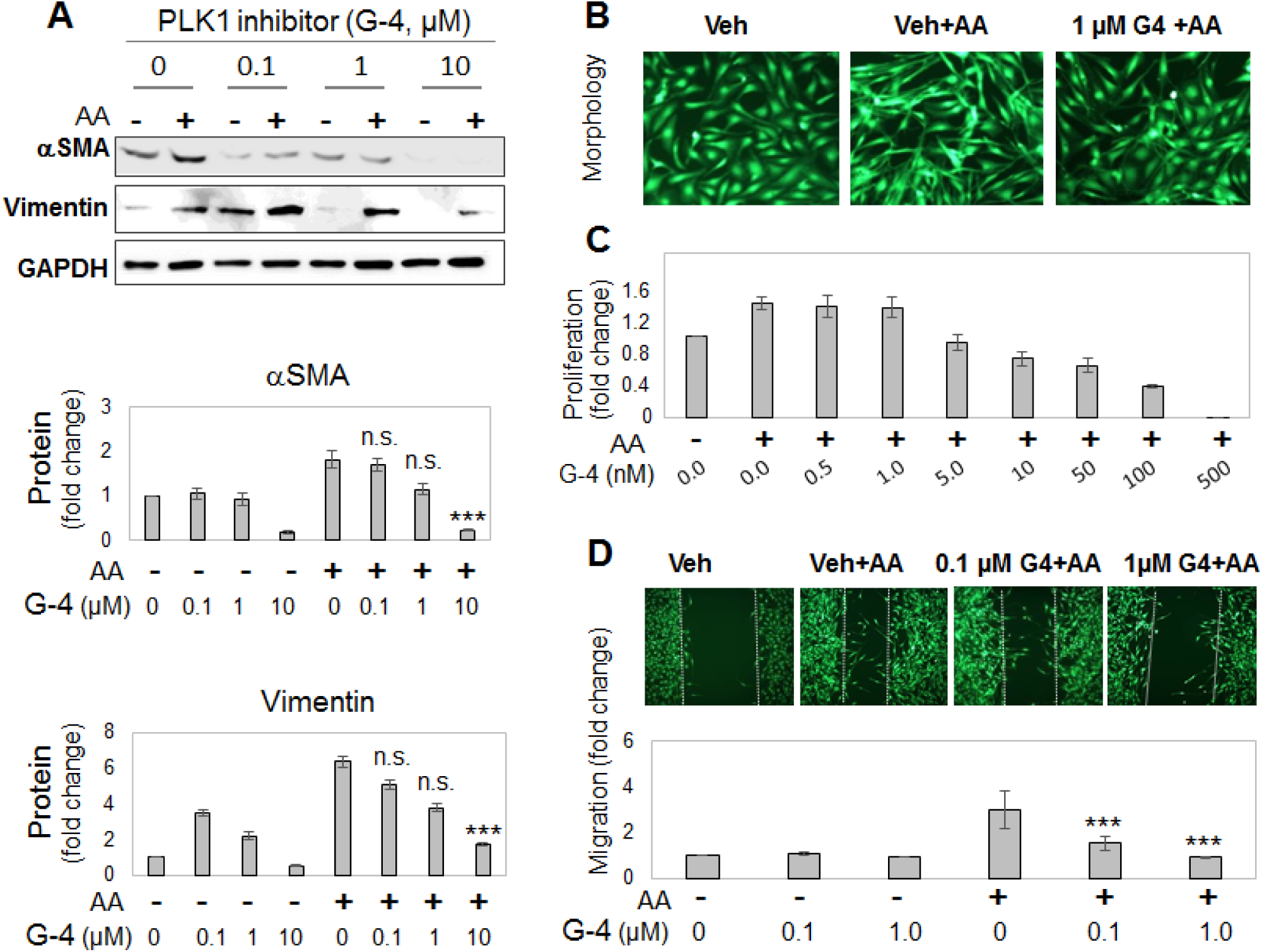
PLK1 inhibition blocks PDGF-AA stimulated fibroblast cell-type transition. Experiments were performed as described in Figure 1 except that the PLK1-selective inhibitor GSK461364 (abbreviated as G-4) was used for pretreatment prior to AA stimulation. A. Western blots of αSMA and vimentin. B. Morphologic comparison. Cells were (or not) stimulated by AA for 24h without or with pre-treatment (1µM G-4). C. Proliferation. CellTiterGlo assay was performed after 72h AA stimulation of cells without or with pretreatment by G-4 at increasing concentrations. D. Migration (scratch assay). Cells were (or not) stimulated by AA for 24h without or with pre-treatment (0.1 or 1 µM G-4). Quantification for A and D: Mean ± SEM, n ≥3 experiments. One-way ANOVA/Bonferroni test: ***P< 0.001 or n.s. (not significant) compared to the condition of AA without G-4.

### Blocking PDGFR kinase activity abrogates AA-stimulated PLK4 activation

Given the profound effects of blocking PLK4 kinase activity on fibroblast cell-type transition, it is important to understand how the activation and expression of PLK4 are regulated. Currently there is very limited information available on this topic. In our experimental setting, PDGF-AA activates PDGFRα, a well-known “gateway” receptor on the cell surface that, in turn, activates a myriad of intracellular pathways^8, 10^. However, whether PDGFRα regulates PLK4 was not previously known. We addressed this question using a kinase inhibitor (Crenolatib; abbreviated as Crenol) selective to both PDGFRα and PDGFRβ (Ki <10 nM)^14^. (PDGFRα-specific compounds are not yet available.)

AA treatment of rat fibroblasts led to the rapid and brief increase in phosphorylation of PDGFRα (at Y754, commonly seen as an upper band)^14^. Specifically, we observed increased phosphorylation within 5 min that declined to the basal level in 20 min. (See Figure 3A.) AA also stimulated phosphorylation of MEK, ERK, JNK, and P38 with a similar time-course as that seen for PDGFRα, consistent with published reports from other cell types^15, 16^. Interestingly, treatment with AA activated PLK4 (phosphorylation at T170)^17^ and PLK1 (phosphorylation at T210)^18^ in the same time scale as that for PDGFRα (Figure 3A). Of note, phosphorylation of AKT and S6K responded to AA stimulation in a delayed manner (Figure 3A), which may mean that these effects are secondary, indirect signaling events. Importantly, while the PDGFR-selective inhibitor Crenol blocked AA-induced activation of PDGFRα as well as MEK/ERK, JNK, AKT and S6K, validating the inhibitor functionality (Figure 3B), this inhibitor also abrogated AA-stimulated activation of PLK4 and PLK1 (Figure 3, C and D), placing them downstream of PDGFRα signaling. Of note, the phospho-PLK4 band most sensitive to AA and Crenol ran at a high position on the blot. Inasmuch as activated PLK4 dimerizes tightly to stabilize the protein^19^, it is tempting to speculate that a transient oligomer may form before its disassembly and degradation. Indeed, phospho-PLK4-positive bands higher than an expected mobility position have been reported elsewhere as well^20, 21^, and recent crystal structures showed a strand-swapped dimer of dimers of the PLK4 PB3 domain^4^, which is absent in other PLKs.

**Figure 3.**
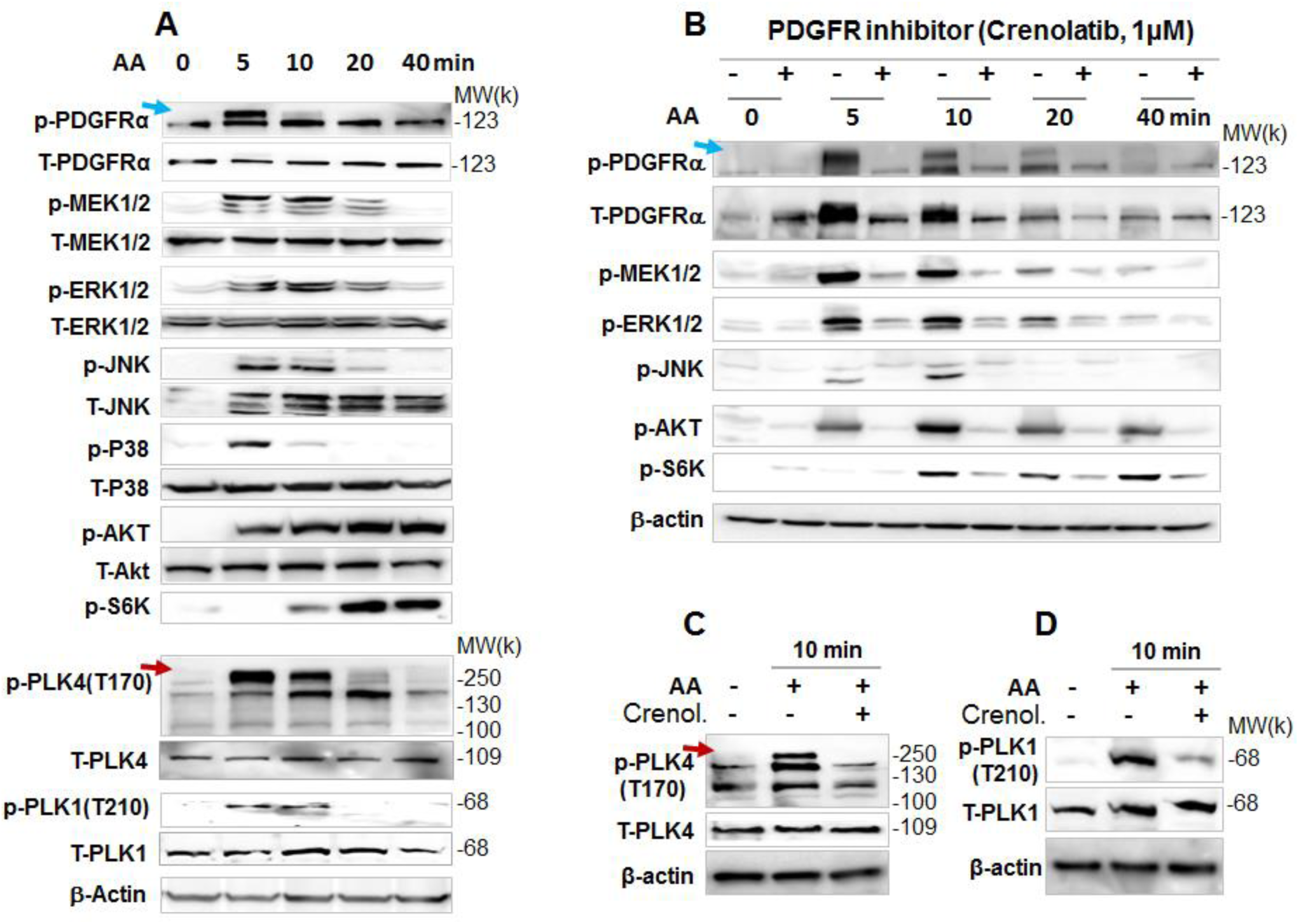
PDGFR inhibition blocks AA-stimulated PLK4 activation. Rat primary adventitial fibroblasts were cultured, starved, and pretreated with vehicle (DMSO) or an inhibitor, as described in Figure 1. Arrows point to the bands of phospho-proteins that were sensitive to both AA stimulation and inhibitor pretreatment. A. Time course of AA-induced activation (phosphorylation) of kinases. Western blots detect a phospho-protein (p-) and its respective total protein (T-). B. Blockade of AA-induced kinase activation by the PDGFR-selective inhibitor Crenolatib (abbreviated as Crenol). Starved cells were pretreated with vehicle or 1 µM Crenol for 30 min prior to AA stimulation. C and D. Blockade of AA-stimulated (10 min) activation of PLK4 (phosphorylation at T170) and PLK1 (phosphorylation at T210) by pretreatment with Crenol (1 µM, 30 min).

### PLK4 activation is regulated by PDGFR downstream kinase activity but not vice versa

To delineate the position of PLK4 in the signaling cascade downstream of PDGFR, we first tested whether its blockade affects the activation of the MAPK and AKT pathways as they were previously implicated as being activated by PDGFR^16^. Our data showed that pretreatment with either the PLK4 inhibitor (Cen-B) or the PLK1 inhibitor (G-4) did not appreciably alter AA-induced phosphorylation of the MAPK pathway kinases (MEK/ERK, JNK, P38) or AKT within 40 min (Figure 4, A and B). However, PLK1 (but not PLK4) inhibition abolished AA-initiated S6K phosphorylation (Figure 4B), suggesting differential functions of these two PLKs. We then dissected which of the PDGFR downstream kinase(s) regulated PLK4 activation, by using a panel of their respective inhibitors. Interestingly, the P38 inhibitor markedly reduced AA-stimulated PLK4 phosphorylation whereas the other inhibitors did not produce a significant effect (Figure 4C). By contrast, PLK1 phosphorylation was significantly attenuated by inhibitors of mTOR, P38, and MEK (Figure 4D). To confirm the specific role of P38 in PLK4 activation, we performed P38 silencing experiments (Figure 4E). The data indicated that PLK4 phosphorylation (10 min after adding AA) was substantially reduced in the cells treated with P38-specific siRNA compared to scrambled siRNA control. Efficient P38 knockdown was indicated by Western blot analysis (Figure 4E, T-P38).

**Figure 4.**
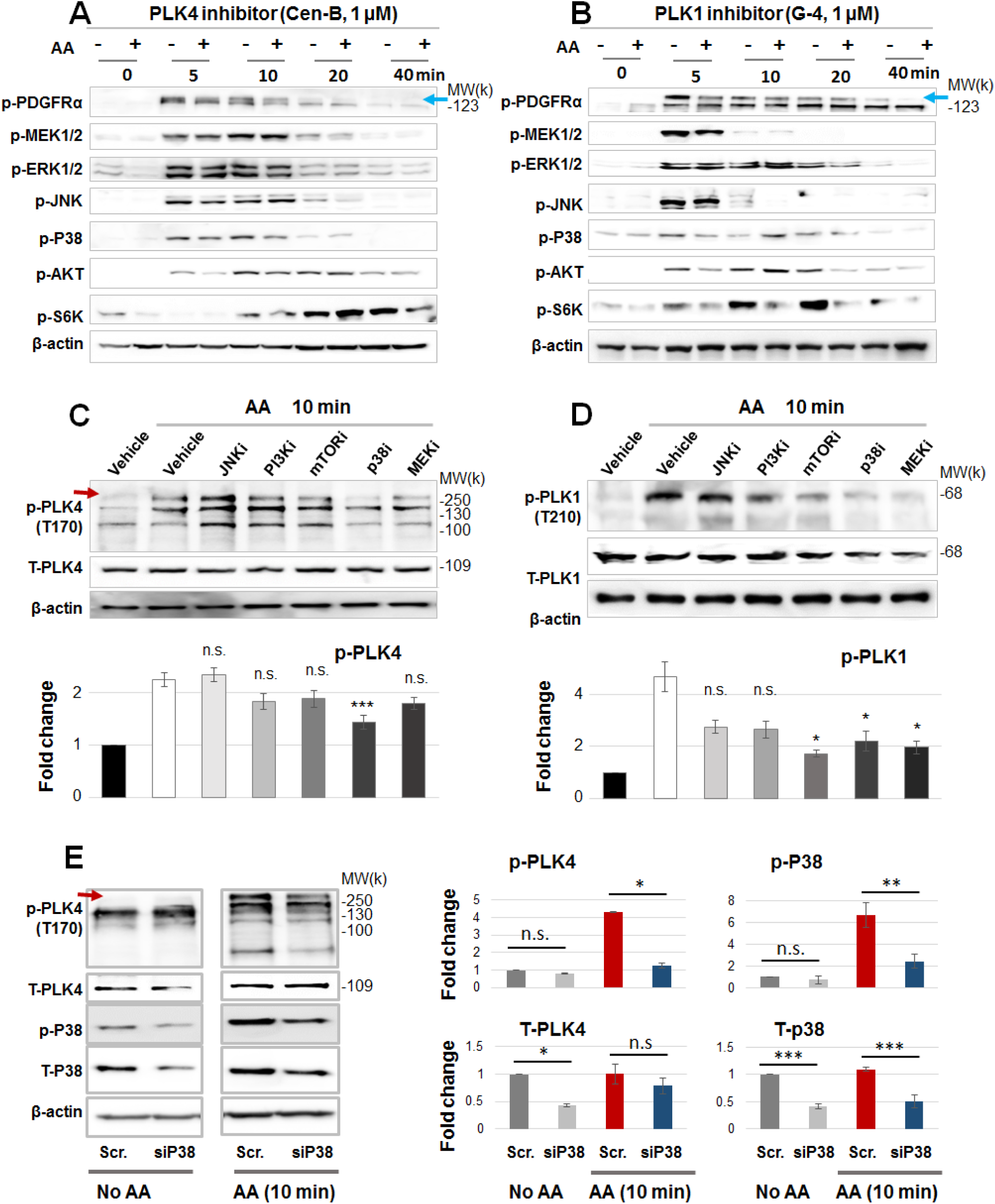
PDGFR downstream regulators of PLK4 activation. A and B. Lack of effect of PLK4 (or PLK1) inhibition on the activation of PDGFRα and downstream kinases. Time-course experiments were performed as described in Figure 3, except that 1 µM Cen-B (PLK4 inhibitor) or 1 µM G-4 (PLK1 inhibitor) were used for a 30-min pre-treatment prior to AA stimulation. C and D. Effects of various kinase inhibitors on the activation of PLK4 and PLK1. Cells were pretreated with an inhibitor of JNK (SP600125), PI3K (LY294002), mTOR (rapamycin), P38 (SB230580), or MEK1/2 (PD98059) for 30 min at 5 µM prior to a 10-min AA stimulation. Quantification: Mean ± SEM, n = 4 or 5 experiments (C or D). One-way ANOVA/Turkey test: *P< 0.05, ***P< 0.001, or n.s. (not significant) compared to the condition of AA without an inhibitor. E. Effect of P38 silencing on the activation of PLK4. Cells were transfected with scrambled or P38α-specific siRNA for 48 h in the complete medium, starved overnight, and then stimulated with AA for 10 min before cell harvest for Western blotting. Quantification: Mean ± SEM, n = 3 experiments. One-way ANOVA/Bonferroni test, *P< 0.05, **P< 0.01, ***P< 0.001, or n.s. (not significant) compared to scrambled siRNA control.

### Blocking PDGFR abrogates AA-stimulated PLK4 protein production

To investigate the long-term regulation of PLK4, we next determined whether PDGFR regulates PLK4 protein levels. As shown in Figure 5A, pretreatment with the PDGFR blocker Crenol concentration-dependently inhibited PLK4 and PLK1 protein upregulation (by AA). Serving as positive control, PDGFR inhibition markedly reduced the protein levels of MEK, ERK, JNK, and P38. In studies to dissect the PDGFR downstream pathways that possibly regulated PLK4 protein expression, we found that pretreatment with inhibitors of these kinases differentially altered PLK4 and PLK1 protein levels (Figure 5, B-F). PLK1 was sensitive to essentially all inhibitors to varied extents. By contrast, PLK4 was sensitive to the P38 inhibitor (Figure 5B), but not to other inhibitors (Figure 5, C-F, at 1 µM), consistent with the results observed on its phosphorylation (Figure 4C).

**Figure 5.**
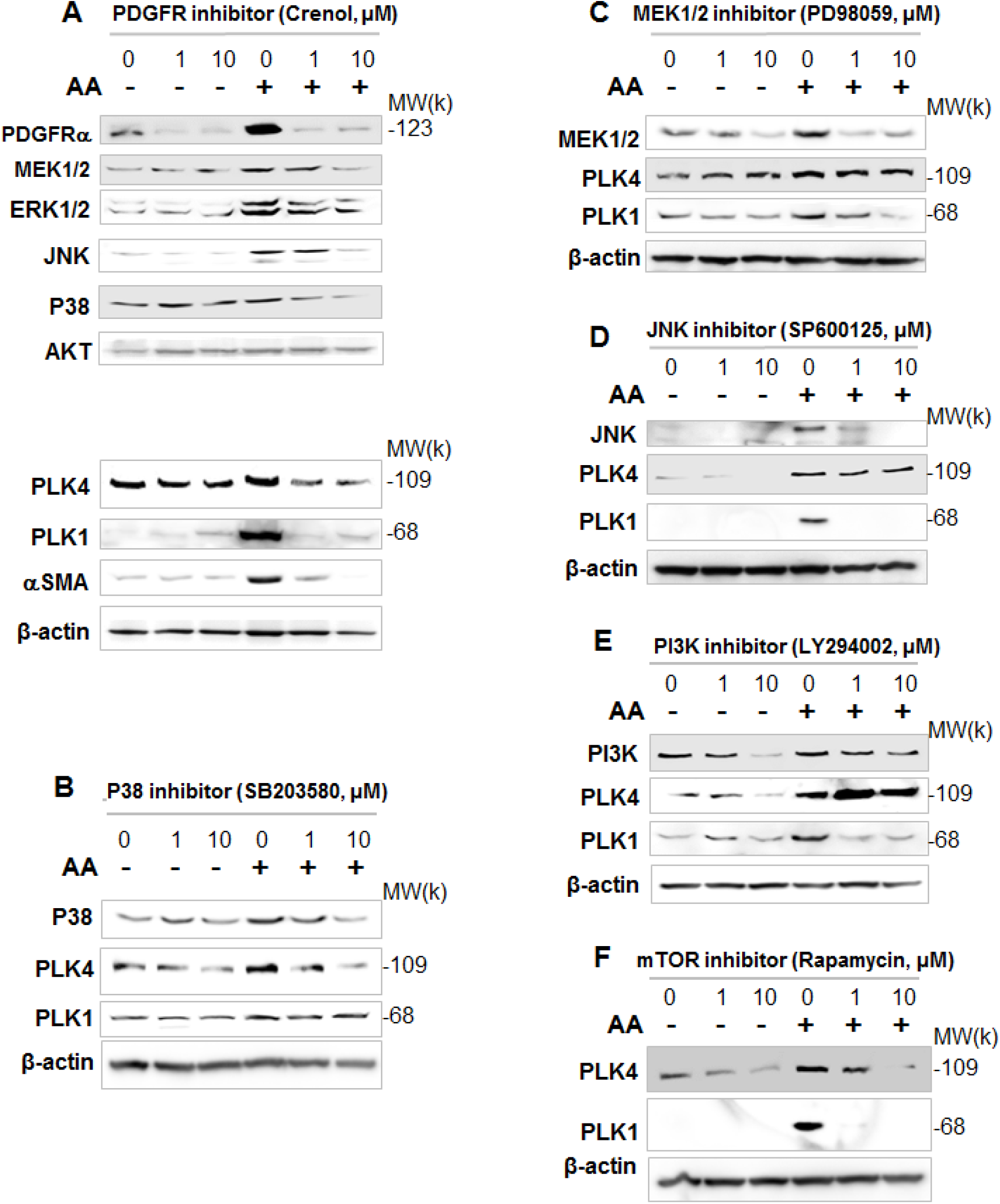
Regulation of PLK4 protein levels by PDGFR and downstream kinases. Rat adventitial fibroblasts were cultured and starved as described in Figure 1. Cells were pre-treated for 30 min with vehicle or an inhibitor for PDGFR (A), P38 (B), MEK1/2 (C), JNK (D), PI3K (E), or mTOR (F) at indicated concentrations, stimulated for 24h with AA, and then harvested for Western blot analysis. Shown are representative blots from two or three similar repeat experiments.

Thus, our results have provided the novel finding that PDGFRs (most likely PDGFRα) stimulates both short-term PLK4 and PLK1 activation and more sustained activity via increased production of these proteins.

### Silencing FoxM1 does not repress the expression of PLK4

We next investigated the transcriptional regulation of PLK4. FoxM1 is a well-known transcriptional activator of mitotic regulatory genes, including cyclin B1, Topo2, Aurora B kinase, and also PLK1^6^. It was therefore logical to ask whether FoxM1 also targets the PLK4 gene. Knocking down FoxM1 with an siRNA, we saw efficient FoxM1 reduction at both the mRNA and protein levels. This FoxM1 reduction led to substantial reduction of PLK1 mRNA and protein expression (Figure 6), consistent with previous reports^5, 6^. However, FoxM1 knockdown increased PLK4 mRNA and did not reduce PLK4 protein levels. Thus, these results implicate differential transcriptional regulations of PLK4 and PLK1.

**Figure 6.**
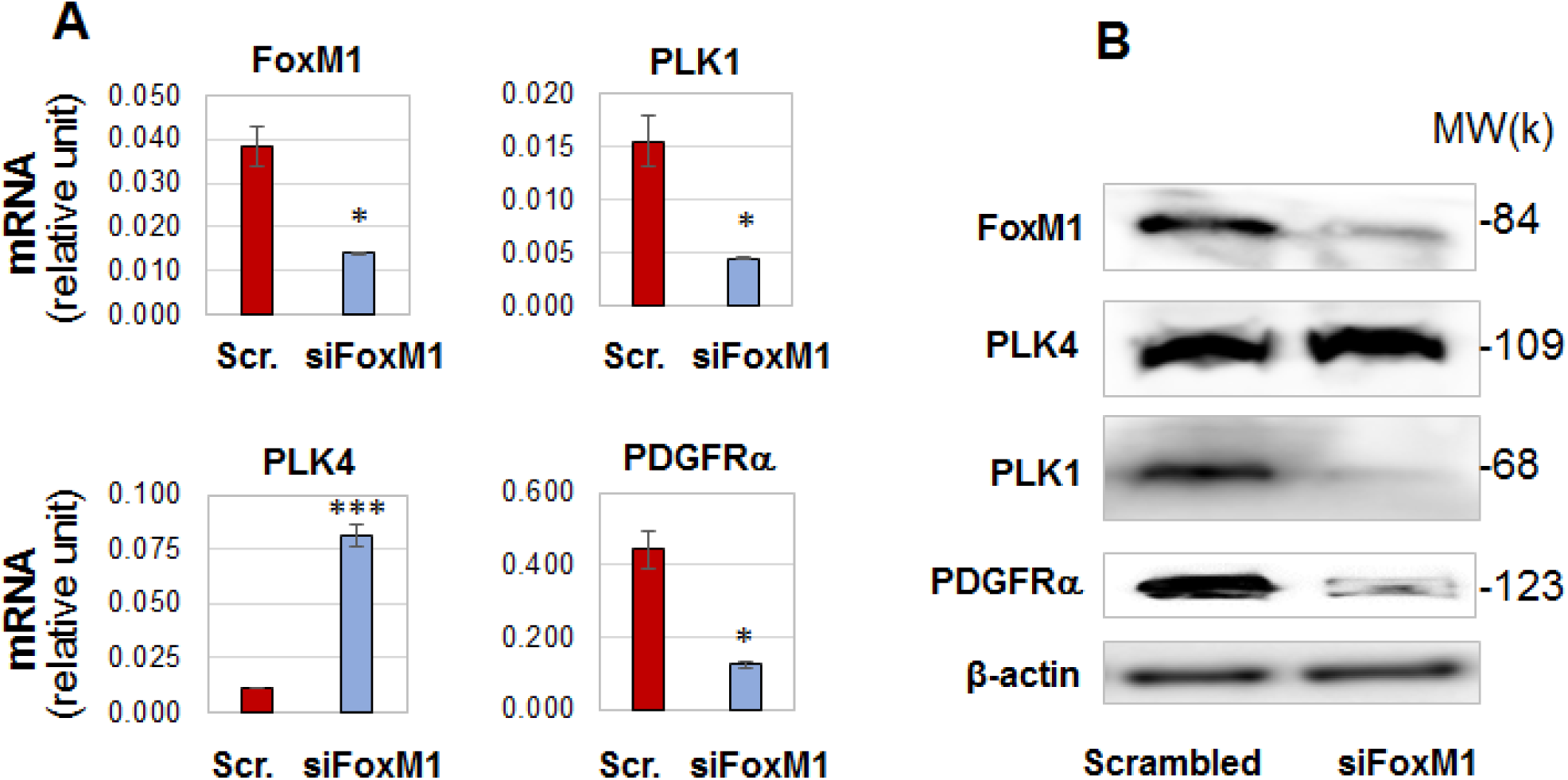
Differential effects of FoxM1 silencing on the transcription of PLK4 and PLK1. Rat adventitial fibroblasts were transfected with scrambled or FoxM1-specific siRNA for 48h and then collected for qRT-PCR (A) and Western blot (B) assays. Quantification: Mean ± SD of triplicate. Student’s t-test: *P<0.05, ***P< 0.001. Shown (in A or B) is one of two similar experiments.

### Pan-BETs inhibition blocks AA-stimulated cell-type transition of rat aortic adventitial fibroblasts

To further identify key regulatory mechanisms that control PLK4 transcription, we explored the role of BETs (bromodomain/extraterminal proteins). While cell identities are governed by specific transcription programs, recent research discovered that BETs function as master epigenetic regulators, directing transcription programs in a cell-type and environment dependent manner^22–25^. Moreover, recent evidence supports an important role for BETs in activation of fibroblasts^26, 27^ albeit without any data obtained from vascular fibroblasts. As shown in Figure 7 (A-D), pretreatment with JQ1 (the first-in-class inhibitor selective for BETs)^28^ abrogated AA-induced myofibroblastic phenotypes, including αSMA and collagen expression, cell migration and proliferation, and also pro-inflammatory cytokines (Figure S4). Apparently, pan-BETs inhibition with JQ1 “phenocopied” the effect of PLK4 inhibition with Cen-B (Figure 1). We were therefore motivated to determine the effect of BETs inhibition on PLK4 (and PLK1) expression.

**Figure 7.**
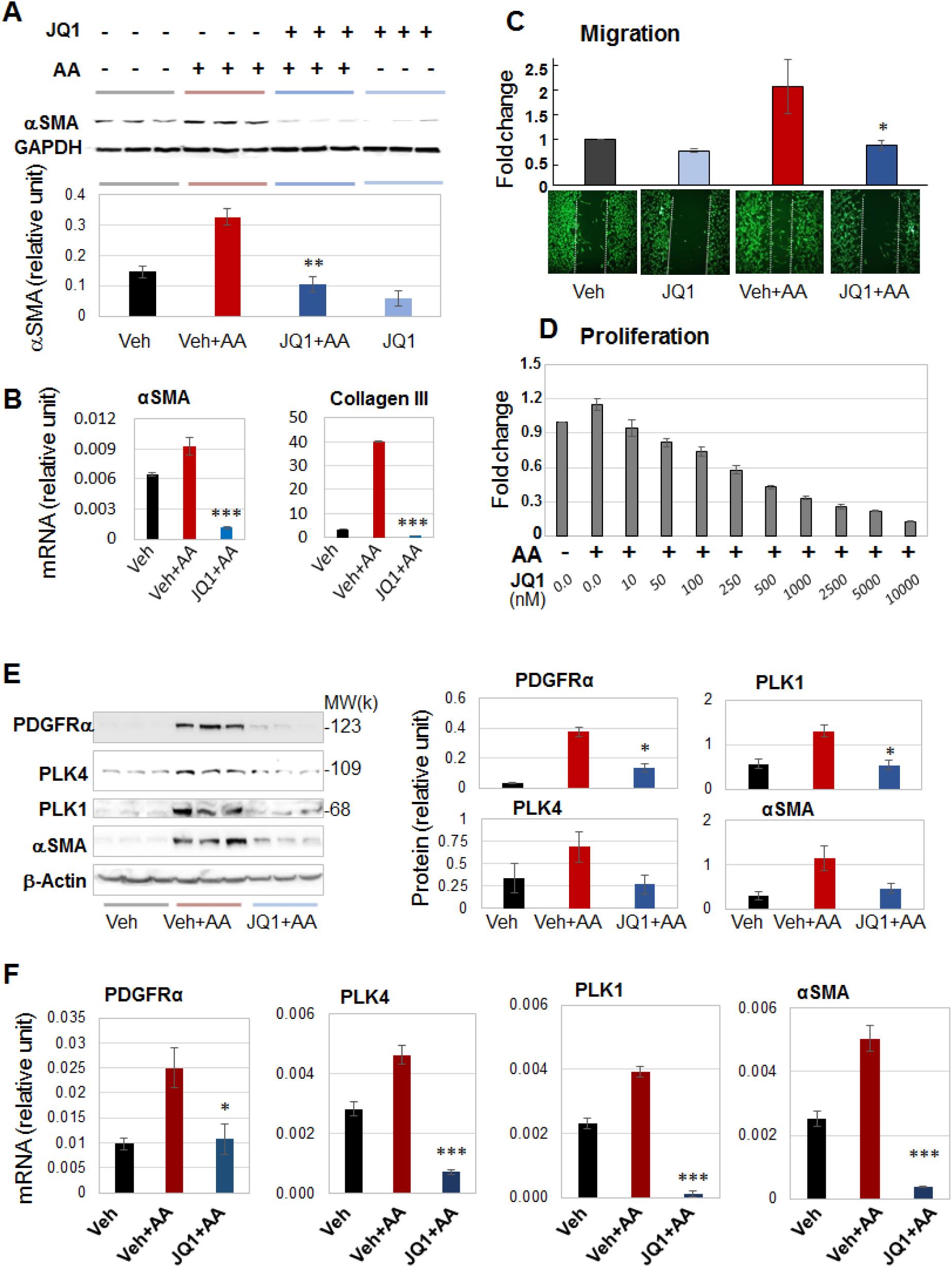
BETs inhibition blocks AA-stimulated fibroblast cell-type transition and expression of PLK4 and PDGFRα. Rat adventitial fibroblasts were cultured and starved as described in Figure 1. Cells were pre-treated with vehicle control or JQ1 (1 µM or otherwise specified) overnight before AA stimulation and cell harvest for various assays. A and B. Western blots and qRT-PCR showing JQ1 blockade of AA-stimulated αSMA expression. Densitometry was normalized to GAPDH. C. Migration (scratch assay). Cells were (or not) stimulated by AA for 48h without or with pre-treatment (1 µM JQ1). Calcein was used to illuminate the cells. D. Proliferation. CellTiterGlo assay was performed after 72h AA stimulation of cells without or with pretreatment by increasing concentrations of JQ1. E. Western blots indicating that pretreatment with JQ1 (1 µM, 2h) abrogated AA-stimulated (24h) protein production of PDGFRα, PLK4, PLK1, and αSMA. Shown are representative blots from one of two similar repeat experiments. Densitometry was normalized to β-actin. F. qRT-PCR data indicating that pretreatment with JQ1 (1 µM, 2h) abrogated AA-stimulated (24h) mRNA expression of PDGFRα, PLK4, PLK1, and αSMA. Quantification (A-F): Mean ± SD of triplicates; shown is one of two similar experiments. One-way ANOVA/Bonferroni test: *P< 0.05, **P<0.01, ***P< 0.001 compared to the condition of AA+vehicle.

### BRD4 predominantly controls the transcription of PLK4 and PDGFRα

Interestingly, pretreatment of the rat fibroblasts with 1 μM JQ1 averted AA-induced upregulation of the mRNAs and proteins of PLK4 and also PLK1 (Figure 7, E and F). Moreover, AA-stimulated PDGFRα mRNA/protein expression was also effectively reduced by pretreatment with JQ1.

The BET family has four known members: BRD2, BRD3, BRD4, and BRD-T (testis-specific and thus irrelevant here). The inhibitor JQ1 binds to all of the BETs^28^. To dissect out which BET was responsible for the potent effect of the pan-BETs inhibitor (JQ1), we performed genetic silencing experiments using siRNAs (Figure 8). Each of the siRNAs gave strong knockdown for its specific target. (See Figure S5.) Whereas silencing BRD4 substantially reduced protein levels of PLK4, PLK1, PDGFRα, and αSMA, silencing BRD2 slightly reduced the protein level of PLK4 but not that of PLK1, PDGFRα, and αSMA; silencing BRD3 did not reduce but appeared to slightly increase the levels of these proteins (Figure 8A). The robust inhibitory effect of silencing BRD4 on PLK4 and other three proteins is indicated by the quantitated data in Figure 8B. Silencing either BRD2 or BRD4 reduced the mRNA levels of PLK4, PLK1, and PDGFRα (though to lesser extents with siBRD2) (Figure 8, C and D). Given these results, BRD4 appeared to be the predominant BET governing the gene transcription of the two PLKs and PDGFRα.

**Figure 8.**
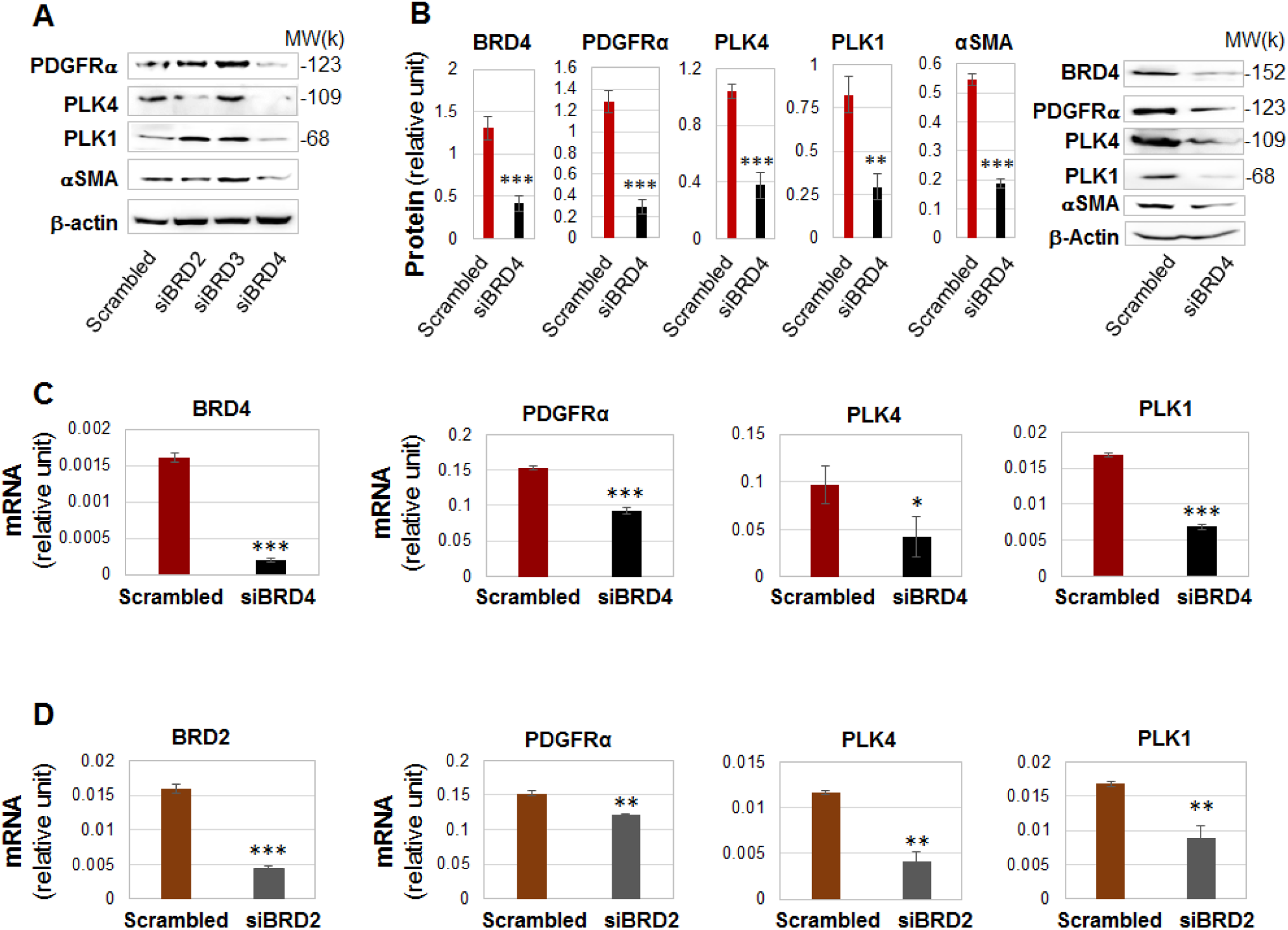
Silencing BRD4 down-regulates the transcription of PLK4 and PDGFRα. Rat adventitial fibroblasts were transfected with a scrambled or BRD-specific siRNA for 48h and collected for Western blot (A) and qRT-PCR (B) assays. A. Comparison of the effects of silencing BRD2, BRD3, or BRD4 on PLK4 protein levels. Shown are representative blots from one of two similar repeat experiments. B. Silencing BRD4 reduces the proteins of PDGFRα, PLK4, PLK1, and αSMA. Quantification: Densitometry normalized to β-actin; mean ± SEM; n = 5 independent experiments, representative blots from one of which are shown on the right. Student’s t-test: **P<0.01, ***P<0.001. C and D. Effects of BRD4 or BRD2 silencing on mRNA levels of PDGFRα, PLK4, and PLK1 (qRT-PCR data). Quantification: Mean ± SD of triplicates; shown is one of two similar experiments. Student’s t-test: *P<0.05, **P<0.01, ***P<0.001.

### Periadventitial administration of PLK4 inhibitor ameliorates vascular fibrosis in the rat carotid artery injury model

After identifying the role for PLK4 in fibroblast cell-type transition and the regulators of its activation and expression, we then determined whether PLK4 inhibition helps reduce fibrosis in vivo. We used a well-established rat arterial injury model in which injury-induced fibrosis manifests as high collagen content in the adventitia. Balloon angioplasty was performed in rat common carotid arteries, and the PLK4 inhibitor (Cen-B) or vehicle control was administered in thermo-sensitive hydrogel distributed around the injured artery, following our previously reported method^24^. Cross-sections of injured arteries collected at day 7 were stained using the Masson’s trichrome method^29^ (Figure 9, A and B). Quantified data indicate that the collagen content (staining intensity normalized to artery perimeter) was significantly reduced in Cen-B treated arteries compared to vehicle (DMSO) control. Consistently, the adventitia thickness also significantly decreased after Cen-B treatment (Figure 9C). No significant change was observed in neointimal hyperplasia (measured as the neointima/media area ratio)^24^. An interesting question for future investigation is whether Cen-B treatment reduces neointima at day 14, a time point when the neointima thickness is maximized. Nonetheless, the results presented herein indicate that in this model of rat carotid artery injury, periadventitial application of the PLK4 inhibitor Cen-B was able to ameliorate vascular fibrosis.

**Figure 9.**
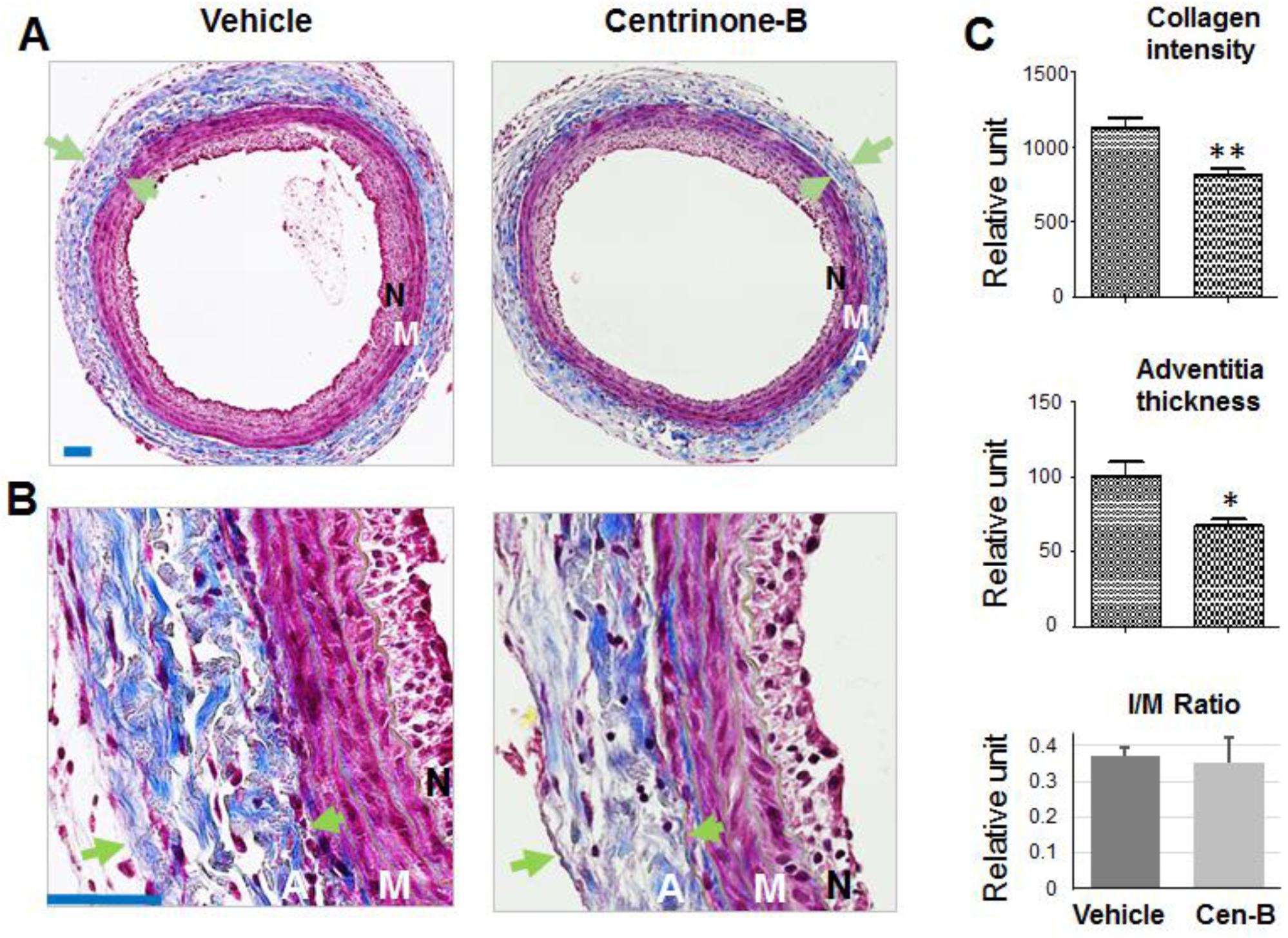
Perivascular administration of PLK4 inhibitor reduces collagen content and thickness of the adventitia in the model of rat carotid artery injury. Following balloon angioplasty of the rat common carotid artery, the PLK4 inhibitor (Cen-B, 100 µg per rat) dissolved in a hydrogel mix was applied around the adventitia of the injured artery. Arteries were harvested at day 7 post injury; cross sections were stained using the Masson’s Trichrome method. A. Representative sections from the arteries treated with vehicle (equal amount of DMSO) or Cen-B. Collagen is stained blue; the adventitia thickness is indicated by two arrows. The anatomy of the artery wall is labeled as A (adventitia), M (media), and N (neointima). B. Magnified areas of the images in A. C. Quantification. Collagen content (staining intensity) and thickness of the adventitia was normalized to the overall vessel size measured as the length of the external elastic lamina (the border between blue and red layers). Neointimal hyperplasia was measured as the intima/media area ratio (I/M). Mean ± SEM, n = 5 animals per group. Student’s t-test: *P<0.05, **P<0.01.

## Discussion

Our major findings are as follows: 1) Though previously known as centriole-specific, PLK4 positively regulated SRF nuclear activity and the target gene αSMA’s transcription. 2) Upon AA stimulation, PLK4 was activated by PDGFR and downstream kinase P38. 3) The transcription of PLK4 was predominantly governed by the epigenetic reader BRD4. Thus, our results revealed a non-canonical role for PLK4 in promoting SRF nuclear activity and fibroblast cell-type transition, and further uncovered a BRD4/PDGFR-dominated mechanism underlying PLK4 expression/ activation. Importantly, PLK4 inhibition was effective in attenuating vascular fibrosis in an arterial injury rat model.

The PLK4’s canonical role in centriole duplication has been recently reviewed^31^. Briefly, PLK4 phosphorylates and complexes with STIL, which in turn recruits SAS6, forming the core module of centriole genesis. However, little has been known about the non-canonical PLK4 function^3, 32^. Thus far, few PLK4’s substrates beyond centriolar proteins have been identified^32^. Thus, PLK4 is much less understood in contrast to PLK1, the best studied PLK member that mediates multiple mitotic processes^1, 31^.

Given that PLK4 is a mitotic factor, it was not surprising to learn that PLK4 is pro-proliferative^1^ in vascular fibroblasts as observed herein, and in cancer progression as previously reported^33, 34^. PLK4 was also recently linked to cancer cell migration and invasion^32^, consistent with our result that PLK4 promoted vascular fibroblast migration. On the other hand, cell-type transition is a process beyond cell proliferation and migration; it involves remodeling of signaling and transcription programs^8^ and thereby results in altered cell type with multiple phenotypic changes. In this regard, it is an unanticipated finding that PLK4, known as a centriole-specific actor, assumes a non-canonical function enabling profound cell-type transformation.

Elevated expression of αSMA endows cells with a contractile function that is governed by the master transcription factor SRF^8^. Consistent with this function, our data showed that, while PLK4 inhibitor profoundly reduced SRF’s activity, PLK4 silencing substantially repressed αSMA levels. This result is somewhat unexpected, given that PLK4, in its centriole-associated function, acts in the cytosol whereas SRF is a chromatin-associated nuclear protein. Interestingly, this paradox may be at least partially explained by our novel observation that PLK4 promotes protein levels (or stability) of MRTF-A. This co-factor of SRF is a cytosol-nucleus shuttling protein and may thereby convey PLK4’s cytosolic function to the nucleus where SRF activation and αSMA transcription occur.

This non-canonical function of PLK4 in SRF activation is intriguing, particularly given that the SRF/αSMA transcriptional remodeling is both phenotypically and mechanistically distinct from cell proliferation and migration, for which evidence of PLK4 regulation can be found in the literature. For example, in a recent report in cancer research PLK4 was found to play a role in cytoskeleton re-organization and lamellipodia formation in HeLa cells^32^, and the authors used the result to explain the PLK4 function in cancer cell migration. Similarly, an earlier study showed that PLK4 expression enhanced the polarity, spreading, and invasion of colon cancer cells^35^. However, whether PLK4 played a role in αSMA transcription was not examined in either of these studies. Another line of evidence is from a very recent report that PLK4 upregulation promoted epithelial cell state transition in neuroblastoma^36^. Although this report showed that PLK4 enhanced vimentin expression, there was no data on αSMA transcription nor on SRF activity. In addition, a new report demonstrated that PLK1 positively regulated angiotensin-II activation of RhoA and actomyosin dynamics in murine vascular smooth muscle cells^11^. However, caution must be taken into the analogy of PLK4 with PLK1. Published evidence and our own results indicate that though in the same family, PLK4 and PLK1 functionally vary, especially in different cell types or states. Nevertheless, the positive regulation of MRTF-A/SRF by PLK4 represents a distinct mechanism that may or may not cross-talk with those previously reported. In this light, a PLK4 non-canonical role in regulating MRTF-A and SRF nuclear activities warrants future studies for further elaboration.

Given PLK4’s critical role in centrosome biogenesis and its potential non-canonical functions, this kinase is expected to be intricately regulated^37^, yet relatively little is known about the regulation of PLK4 in mammalian systems^17, 38^. We found that PLK4 activation (phosphorylation at T170^17, 37^) by AA treatment was blocked by PDGFR inhibition. Because AA specifically activates the PDGFR αα homodimer^10^, we inferred that it was primarily the activation of PDGFRα that led to PLK4 activation. We reasoned that PDGFRα activation of PLK4 was indirect because PDGFRα is a receptor tyrosine kinase^10^ with no known threonine kinase activity. Downstream of PDGFR, P38 appeared to be an activator of PLK4, as evidenced by experiments using either a P38 inhibitor or siRNA. By contrast, PLK4 inhibition did not affect phosphorylation of PDGFRα or P38, placing PLK4 downstream of the PDGFR/ P38 signaling. Our results do not reveal whether PLK4 is a direct substrate of the P38 kinase, as definitively addressing this question requires an assay using both purified proteins. Although a p38-mediated PLK4 activation has not been previously reported, evidence consistent with this finding comes from a recent in vivo study where P38 proved to be a critical myofibroblastic activator^39^. Consistent evidence also comes from our result that inhibiting either PDGFR or P38 averted AA-induced PLK4 protein upregulation. In contrast, pretreatment with inhibitors of other PDGFR downstream pathways (e.g. MEK/ERK and JNK) did not produce an obvious effect on PLK4 protein levels. Taken together, our results have profiled a PDGFR-> P38->PLK4->MRTF-A->SRF signal transduction pathway (Figure 10).

**Figure 10.**
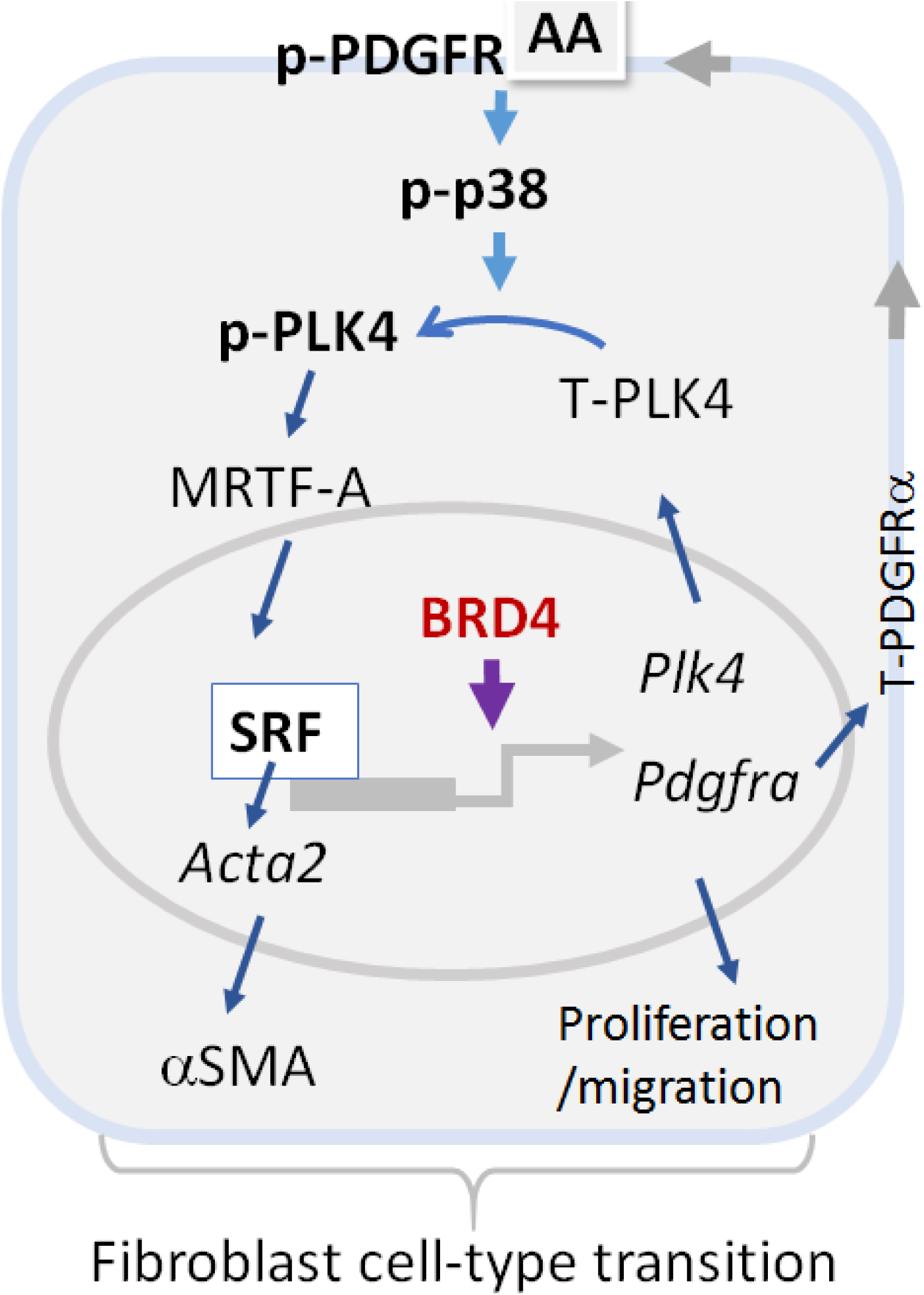
Schematic of the pathways that regulate PLK4 activation and expression. In an acute phase (within 5 min), AA as extracellular signal initiates activation of the PDGFR(α)->P38->PLK4 pathway. PLK4 activation results in elevation of MRTF-A which in turn activates SRF and αSMA expression. At the transcription level, BRD4 as an epigenetic determinant promotes the transcription of PLK4, and also PDGFRα which further enhances PLK4 activation.

In pursuit of the molecular mechanism underlying transcriptional regulation of PLK4, we investigated representatives of two different categories of regulators, namely those known to regulate members of the PLK family (FoxM1) and those known to regulate myofibroblastic activation (BETs). Interestingly, our data showed that silencing the transcription factor FoxM1, a known regulator of PLK1, did not repress PLK4 expression, either at mRNA or protein levels. Instead, we found that BET family member BRD4 played a predominant role in governing PLK4 gene transcription. This is consistent with previous reports indicating a positive role of BETs in fibroblast activation^26, 27^. These studies showed that BET family inhibitors mitigated myofibroblast transdifferentiation and fibrosis in vital organs such as lung, liver, pancreas, and heart^25–27, 40^. Our study differs from these reports by addressing two important questions: 1) Do any of the BETs regulate PDGFRs? 2) Do any of the BETs regulate PLK4 or PLK1? Given our finding that silencing BRD4 effectively reduces mRNA and protein levels of PDGFRα, PLK4, and αSMA, BRD4 appears to play a master role in governing this entire PLK4 pathway. Indeed, in the recent literature BRD4 emerges as a master epigenetic regulator in a variety of cell-state/type transitions^23–26^. While a cell-type/state transition often results from environmental perturbations, environmental cues (e.g. PDGF-AA) and the resultant transcription reprogramming represent two ends of this event while an epigenetic regulator such as BRD4 functions at their interface. Upon stimulation, BRD4 acts as a key organizer of trans-and cis-elements (e.g. super-enhancers) and the core transcription machinery, and (re)localizes them to specific sets of genes to activate their transcription. The two tandem bromodomains of BETs (blocked by JQ1) “usher” this transcription assembly to target genes by binding to bookmarked (acetylated) chromatin loci. This BRD4-directed mechanism has been recently recognized as critical in orchestrating cell-state transitions associated with various pathobiologic contexts^22–26^. As indicated by our data, BRD2 also participated in regulating the PLK4 pathway but played a lesser role (*vs* BRD4). The mechanistic difference between the effects of BRD4 and BRD2 on PLK4 awaits further investigation.

## Conclusions

We have made an unexpected finding that PLK4, a kinase traditionally known as a centriole duplication factor, also regulates cell-type transitions of vascular fibroblasts under PDGF-AA stimulation. The significance of our study is three-fold. First, PLK4’s positive regulation in MRTF-A/SRF activation is a novel non-canonical PLK4 function. Second, while PLK4 is a central player in the PDGFRα-> P38->PLK4->SRF pathway that prompts PDGF-induced αSMA expression, BRD4, as an epigenetic determinant, governs the entire pathway. In this context, PLK4 appears to be a novel signaling effector in sensing and transmitting environmental cues. Third, consistent with the in vitro role of PLK4 in fibroblast cell-type transition, PLK4 inhibition mitigates vascular fibrosis in vivo. Now that clinical tests are ongoing for PLK4 as an anti-cancer therapeutic target^31, 33, 41^, rectifying PLK4 activity may provide a new option for developing anti-fibrotic interventions.

## Methods

### Animals and ethics statement

Male Sprague–Dawley rats were purchased from Charles River Laboratories (Wilmington, MA), housed and fed under standard conditions, and used for in vivo experiments at body weights of 300–330 g. All animal studies conformed to the Guide for the Care and Use of Laboratory Animals (National Institutes of Health) and protocols were approved by the Institutional Animal Care and Use Committee at The Ohio State University. Isoflurane general anesthesia was applied during surgery (through inhaling, at a flow rate of 2 L/minute), and buprenorphine was subcutaneously injected (0.03 mg/kg, ∼0.01 mg/rat) after the surgery. Animals were euthanized in a chamber gradually filled with CO_2_.

### Rat aortic adventitial fibroblast cell culture, induction of cell-type transition, and pretreatment with inhibitors

Primary aortic adventitial fibroblasts were isolated from 6-8 weeks old male Sprague-Dawley rats. For cell expansion, the culture was maintained at 37°C/5% CO_2_ in Complete Fibroblast Medium (Cat. M2267, Cell Biologics Inc.) containing growth factor supplement and 10% fetal bovine serum (FBS, Cat. 6912, Cell Biologics Inc.); 0.25% Trypsin-EDTA solution (Cat. #25200114, Life technologies, Carlsbad, CA.) was used for cell detachment. The fibroblasts at passage 5 were used for experiments. For induction of fibroblast cell-type transition, cells were first starved overnight in Fibroblast Basal Medium (Cat. 2267b, Cell Biologics Inc.) that contains no FBS, and then stimulated with 60 ng/ml PDGF-AA (rat recombinant, R&D Systems Inc., MN) for specifically indicated (in figures) length of time. In the experiments using various inhibitors (Table S1), prior to adding PDGF-AA cells were pretreated with an inhibitor for 2h (at an indicated concentration) or vehicle control (equal volume of DMSO, Sigma-Aldrich, St. Louis, MO).

### Cell proliferation and migration assays

Cell viability assay was performed using the CellTiterGlo kit (Cat. G7571, Promega, Madison WI), as we previously reported^24^. Briefly, 72h after PDGF-AA treatment, plates were decanted and re-filled with 50 μl of CellTiter-Glo reagent and 50 μl PBS per well. Plates were incubated at room temperature for 20 min and then read in FlexStation 3 Benchtop Multi-Mode Microplate Reader (Molecular Devices, Sunnyvale, CA) by using a 250-ms integration.

Cell migration was measured using the scratch assay following our previous report^24^. Briefly, cells were cultured to 90% confluence in 6-well plates and then starved for 24 h in Fibroblast Basal Medium. A sterile pipette tip was used to generate an ∼ 1 mm cell-free gap. Dislodged cells were washed away with PBS. Plates were then refilled with Fibroblast Basal Medium containing 20 ng/ml PDGF-AA and no FBS and incubated for 24 h. For pre-treatment prior to PDGF-AA stimulation, an inhibitor or vehicle control (equal amount of DMSO) was incubated with the cells for 2h or otherwise specified. For illumination of the cells, <u>Calcein</u>-AM was added (final 2 μM) and incubated for 30 min at the end of the PDGF-AA treatment, and images were taken after 3 times of wash with PBS. Cell migration was quantified with ImageJ (NIH) based on the width of the cell-free gap.

### Western blotting to assess protein levels

Western blot analysis was done as we previously described^24^. Briefly, cells were harvested and lysed on ice in the RIPA buffer (Cat.89900, Thermo Fisher) that includes a protease inhibitor cocktail (Cat.87785, Thermo Fisher). Cell lysates were quantified for protein concentrations using the Bio-Rad DC™ Protein Assay kit (Cat.5000112, BioRad) and loaded to a 10% SDS-PAGE gel. The full primary antibody list is shown in Table S2. Beta-actin or GAPDH was used for loading control. The signals from primary antibodies were amplified by horseradish peroxidase (HRP)-conjugated immunoglobulin G (IgG) (Bio-Rad) and illuminated with Pierce ECL Western Blotting Substrates (Thermo Fisher). Blot images were immediately recorded with Azure C600 Imager (Azure Biosystems). Protein band densitometry was quantified using NIH ImageJ and normalized to loading control for statistical analysis.

### Quantitative real-time PCR (qRT-PCR) to determine mRNA levels

We followed the method described in our previous report^24^. Briefly, total RNA was isolated from cultured cells using the Trizol reagent (Invitrogen, Carlsbad, CA). Potential contaminating genomic DNA was removed by using gDNA Eliminator columns provided in the kit. RNA was quantified with a Nanodrop NP-1000 spectrometer, and 1 µg was used for the first-strand cDNA synthesis. Quantitative RT-PCR was then performed using Quant Studio 3 (Applied Biosystems, Carlsbad, CA). The house keeping gene GAPDH was used for normalization. Each cDNA template was amplified in triplicate PerfeCTa SYBR® Green SuperMix (Quantabio) with gene specific primers listed in Table S3.

### Gene silencing with siRNAs

siRNAs were ordered from Thermo Fisher (sequences listed in Table S4). Cells were grown to ∼70% confluence in 6-well plates in Complete Fibroblast Medium. A gene-specific siRNA was added to transfect fibroblast cells for overnight using the RNAi Max reagent (Cat.13778150, Thermo Fisher). Cells then recovered in the complete medium for 24h. For induction of fibroblast cell-type transition, the culture was changed to Fibroblast Basal Medium and incubated for overnight prior to AA stimulation.

### Luciferase reporter assay for SRF transcriptional activity

We followed the manufacturer’s instruction and our recently reported protocol^12^. Briefly, the pGL4.34 vector plasmid containing the CArG box (SRF response element) was purchased from Promega (Cat.E1350). An empty vector was generated by removing the SRF response element. Cells were transfected with the empty vector (control) or pGL4.34 using jetPRIME® Transfection Reagent (Cat.114-07, Polyplus-transfection Inc., NY, USA). Positively transfected cells (HEK293) were selected with hygromycin B (Cat.10687010, Thermo Fisher), seeded in 24-well plates at a density of 20,000 cells/well, and grown for 6 h. Cells were treated with vehicle (DMSO) or 1 µM centrinone-B for 2 h, and then lysed in Bright-Glo (Cat.2610, Promega), and luminescence was read in FlexStation 3 Benchtop Multi-Mode Microplate Reader (Molecular Devices, Sunnyvale, CA).

### Model of rat carotid artery injury and peri-adventitial administration of PLK4 inhibitor

To induce adventitial fibrosis, balloon angioplasty injury was performed in rat common carotid arteries as we previously described^24^. Briefly, rats were anesthetized with isoflurane (5% for inducing and 2.5% for maintaining anesthesia). A longitudinal incision was made in the neck to expose carotid arteries. A 2-F balloon catheter (Edwards Lifesciences, Irvine, CA) was inserted through an arteriotomy on the left external carotid artery and advanced into the common carotid artery. To produce arterial injury, the balloon was inflated at a pressure of 2 atm and withdrawn to the carotid bifurcation and this action was repeated three times. The external carotid artery was then permanently ligated, and blood flow was resumed. Immediately following balloon injury, a PLK4 inhibitor (centrinone-B, 100 µg/rat) or DMSO control dissolved in a mix of two thermosensitive hydrogels was administered around the adventitia of injured arteries. The hydrogel mix (total 400 µl) contained equal volume of 20% AK12 (PolySciTech, AKINA Inc.) and 25% Pluronic gel (Sigma-Aldrich). One week after balloon injury, common carotid arteries were collected from anesthetized animals following perfusion fixation at a physiological pressure of 100 mm Hg. Throughout the surgery, the animal was kept anesthetized via isoflurane inhaling at a flow rate of 2 L/minute. Buprenorphine was subcutaneously injected (0.03 mg/kg, ∼0.01 mg/rat) after the surgery. Animals were euthanized in a chamber gradually filled with CO_2_.

### Morphometric analysis

Cross sections (5 µm thick) were excised from rat common carotid arteries embedded in paraffin blocks. Sections were stained for collagen and morphometric analyses using a Masson’s trichrome approach^29^ with reagents from Abcam (Cat.ab150686). Collagen was stained blue and smooth muscle actin (media layer) was stained red. Fibrosis was assessed by two parameters in parallel: the thickness and collagen content of the adventitia layer which was distinguishable from other tissue layers by distinct colors. Collagen content was measured as blue stain intensity normalized to the artery overall size (length of external elastic lamina). Planimetric parameters for assessing intimal hyperplasia (intima/media area ratio) were measured following our previous report^24^: area inside external elastic lamina (EEL area), area inside internal elastic lamina (IEL area), lumen area, intima area (= IEL area-lumen area), and media area (= EEL area – IEL area). All measurements were performed with ImageJ by a student blinded to treatment groups. The data from all 3-5 sections were pooled to generate the mean for each animal. The means from all the animals in each treatment group were then averaged, and the standard error of the mean (SEM) was calculated.

### Statistical analysis

Differences in measured variables between experimental conditions were assessed by one-way ANOVA with post hoc Bonferroni test or Student’s t-test (comparison between two conditions); p < 0.05 was considered significant. Data are presented as mean ± SEM (from at least 3 independent experiments) or mean ± SD (of triplicates). Statistics and graphical data plots were generated using GraphPad Prism v.5.0 for Windows (GraphPad).

## Acknowledgement

This work was supported by NIH grants R01 HL133665 (to L.-W. G.) and R01HL-143469 (to K.C.K., L.-W. G.), and an AHA pre-doctoral award 17PRE33670865 (to M.X.Z.).

We would like to thank Dr. Jing Zhao for discussion and critical comments.

## Supplemental figures

**Figure S1.**
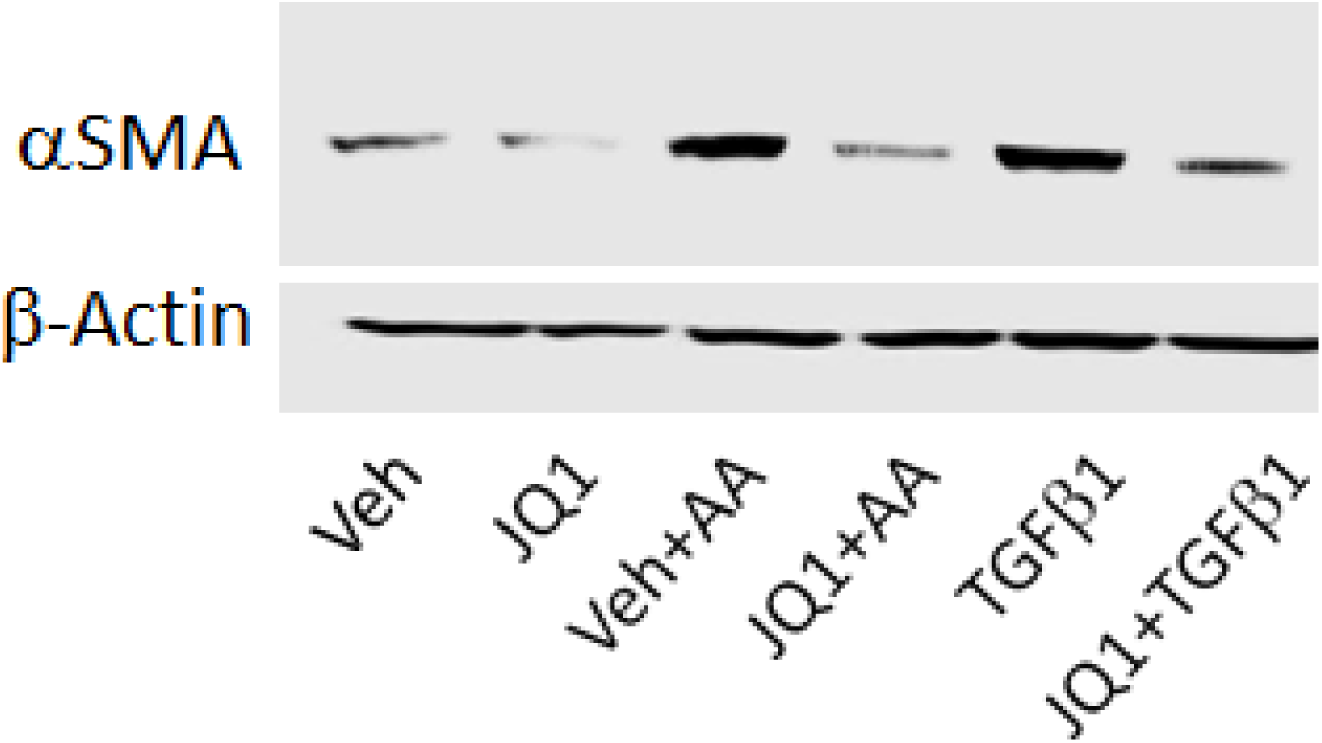
PDGF AA effectively stimulates αSMA expression in rat adventitial fibroblasts. Rat primary adventitial fibroblasts were cultured in the complete medium, starved in the basal medium overnight, and pretreated for 2h with vehicle (equal amount of DMSO) or an inhibitor (JQ1), followed by stimulation with 60 ng/ml PDGF-AA or 20ng/ml TGFbeta. Cells were harvested at 24h after stimulation for Western blot analysis. Shown is one of two similar experiments.

**Figure S2.**
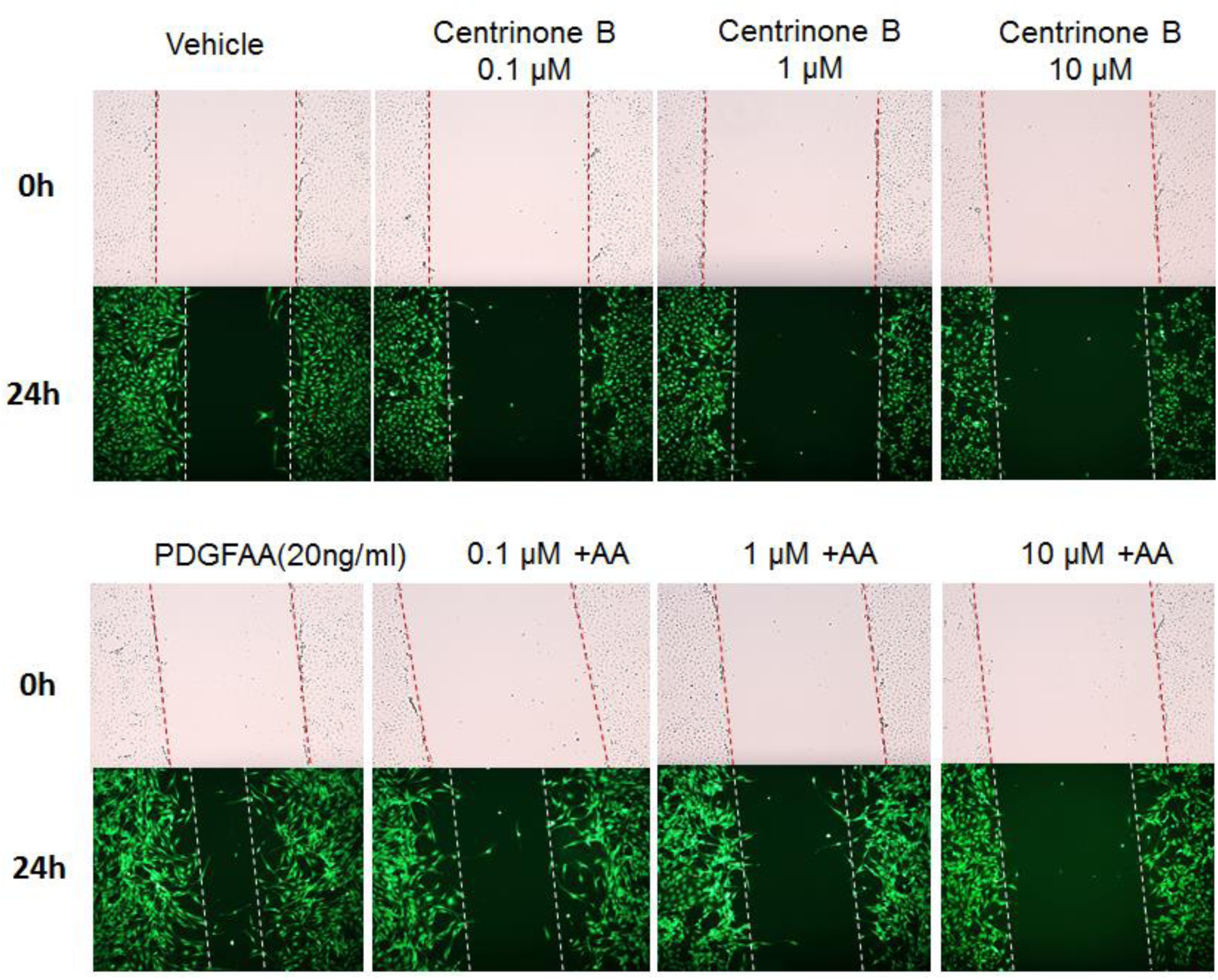
Migration assay in Figure 1. Migration (scratch assay) was performed in the same experiments as in Figure 1. Briefly, rat primary adventitial fibroblasts were cultured, starved, pretreated with inhibitor prior to AA stimulation.

**Figure S3.**
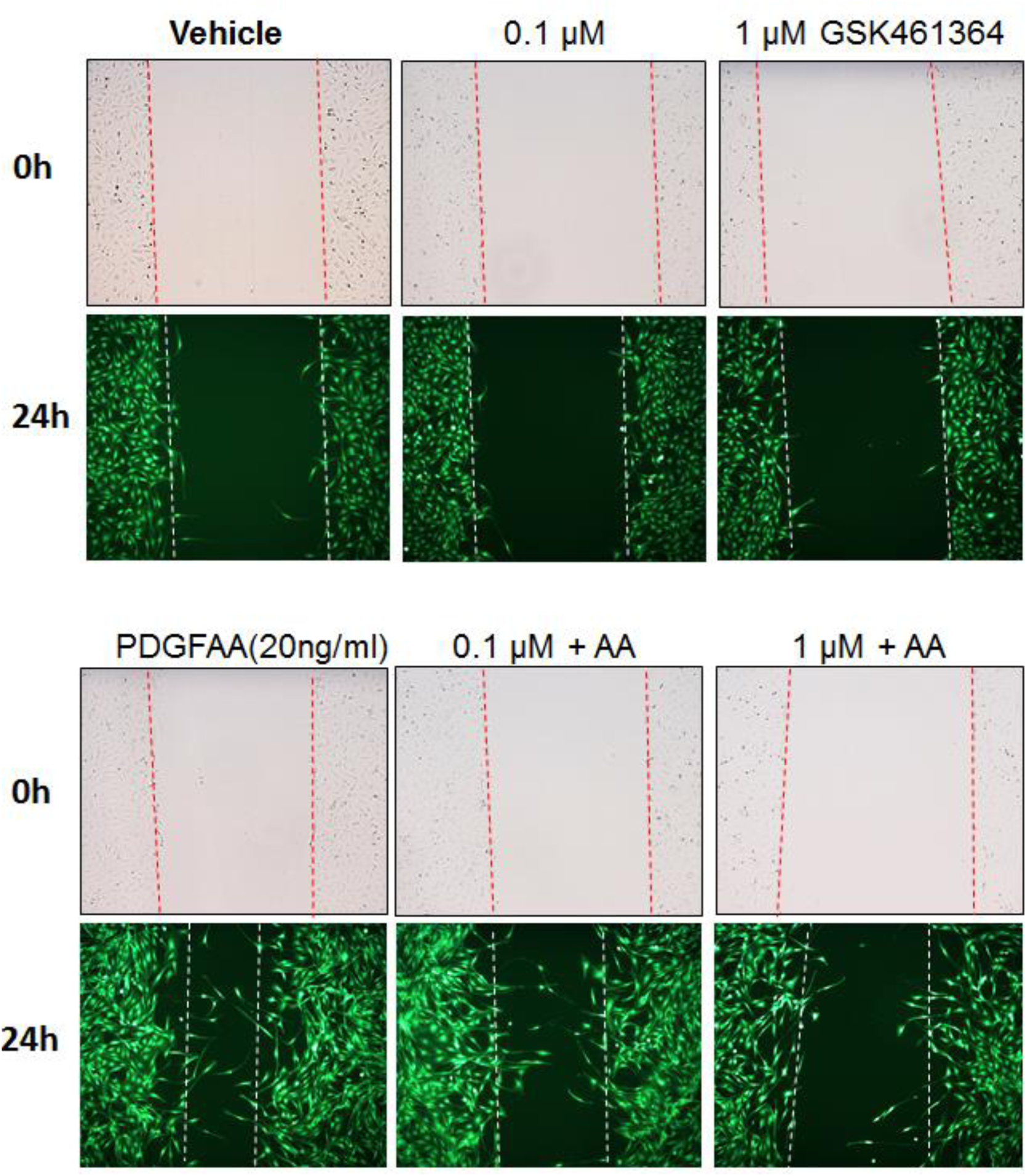
Migration assay in Figure 2. Migration (scratch assay) was performed in the same experiments as in Figure 2. Briefly, rat primary adventitial fibroblasts were cultured, starved, pretreated with inhibitor prior to AA stimulation.

**Figure S4.**
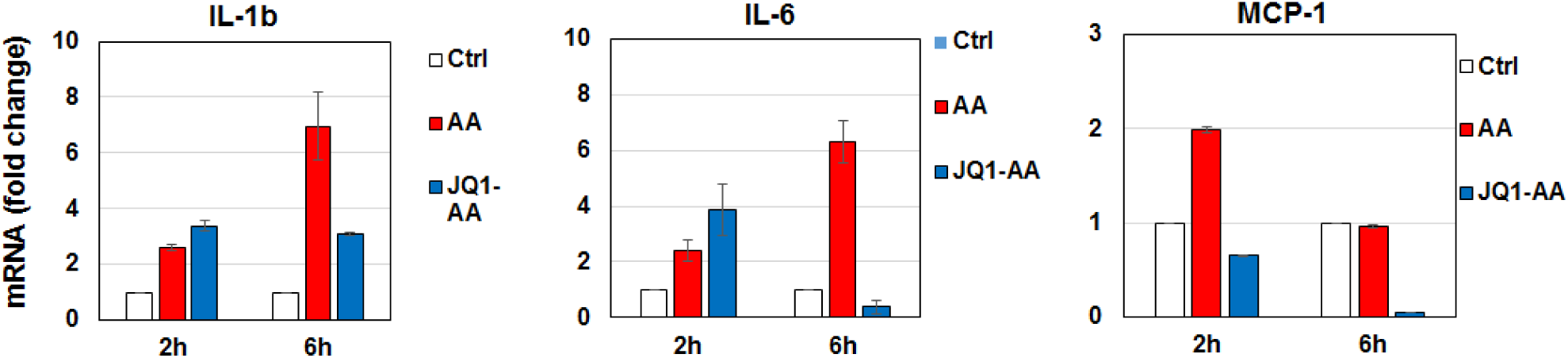
JQ1 inhibits AA-stimulated inflammation and collagen expression. The qRT-PCR assay was performed in the same experiments as in Figure 7. Briefly, rat primary adventitial fibroblasts were cultured, starved, pretreated with JQ1 prior to AA stimulation. Quantification: Mean ± SD, n =3; one of two similar repeat experiments.

**Figure S5.**
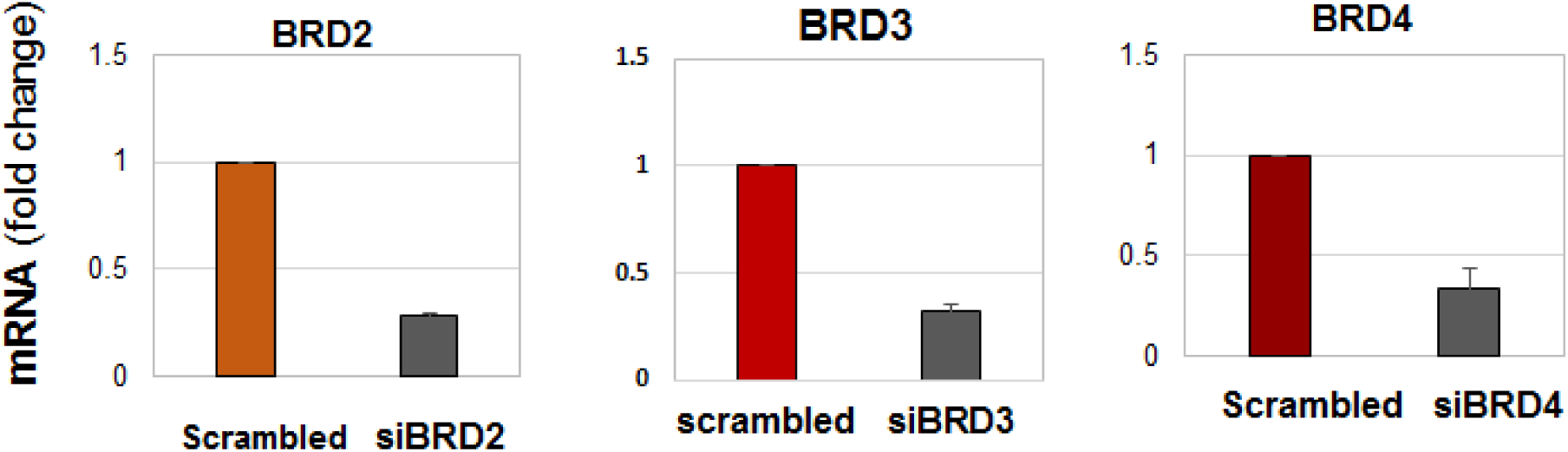
siRNA knockdown efficiency for BRD2, BRD3, and BRD4. The qRT-PCR assay was performed in the same experiments as in Figure 8. Quantification: Mean ± SD, n =3; one of two similar repeat experiments.

## Supplemental Tables

**Table S1.** BET and kinase inhibitors

| Inhibitor | Target protein | Source company | Catalog no. | Stock solution concentration |
| --- | --- | --- | --- | --- |
| JQ1(+) | Bromodomain inhibitor | ApexBio | A1910 | 10mM |
| GSK461364 | PLK1 | ApexBio | A8441 | 10mM |
| Centrinone B | PLK4 | Tocris | 5690 | 10mM |
| Crenolanib | PDGFR | MedChemExpress | HY-13223 | 10mM |
| Tubastatin A | HDAC6 | Selleckchem | 1252003-15-8 | 10mM |
| SB230580 | P38 | Calbiochem | 559389 | 10mM |
| SP600125 | JNK | Calbiochem | 420119 | 20mM |
| LY294002 | PI3K | Calbiochem | 440202 | 20mM |
| Rapamycin | mTOR | Calbiochem | 553210 | 10mM |
| PD98059 | MEK1/2 | Calbiochem | 513000 | 25mM |

**Table S2.**
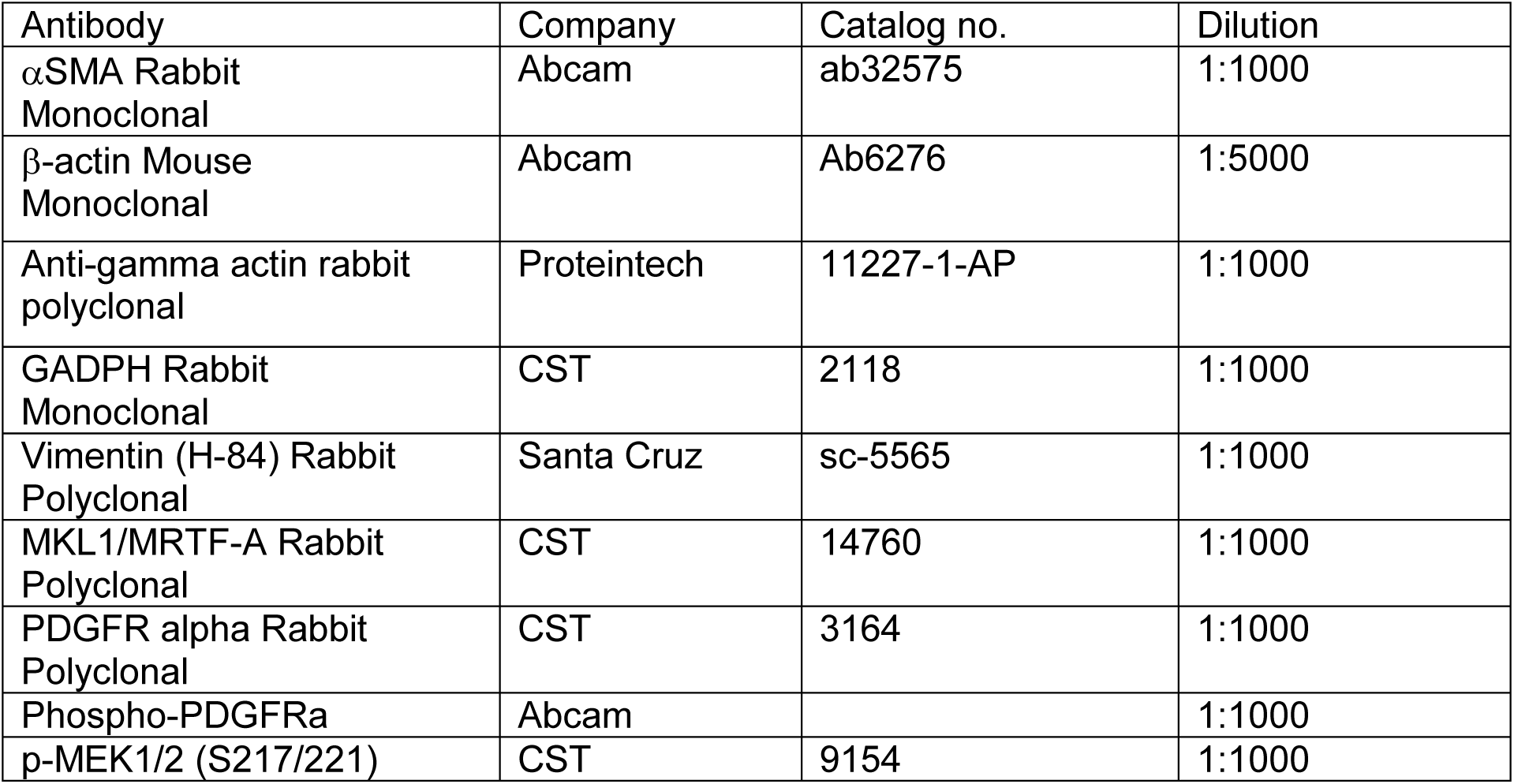
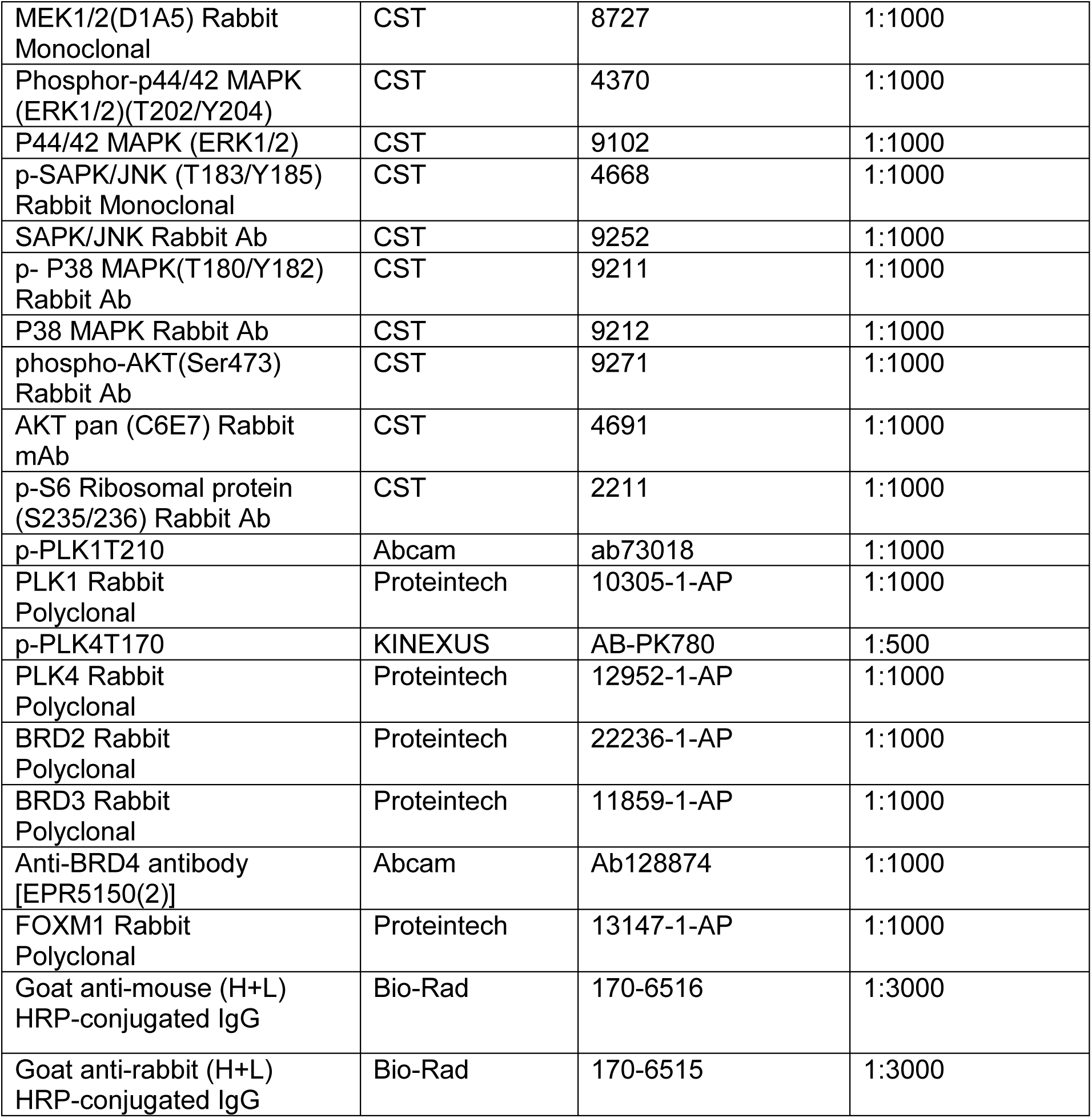
Antibodies for Western blotting

**Table S3.**
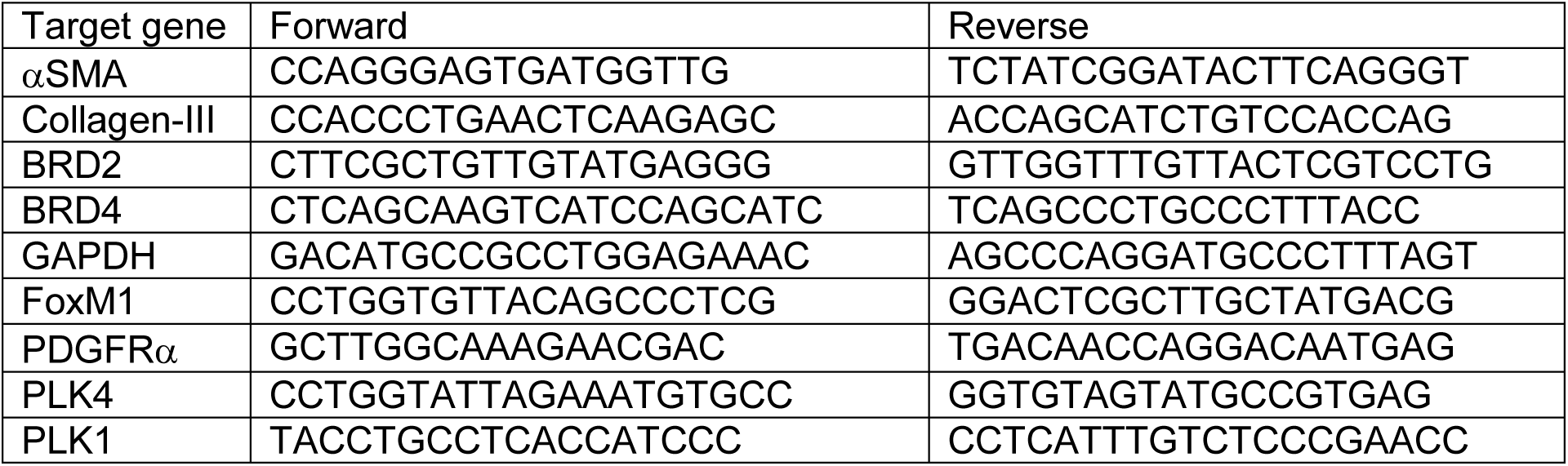
Primers for qRT-PCR (rat sequence)

**Table S4.**
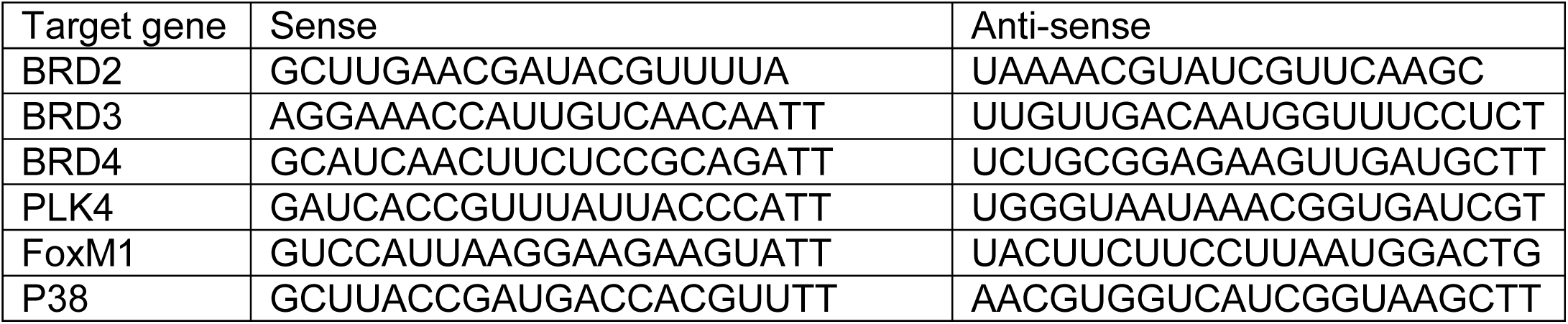
siRNAs (rat sequence)

